# A Suite of Advanced Tutorials for the WESTPA 2.0 Rare-Events Sampling Software [Article v0.1]

**DOI:** 10.1101/2022.10.04.510803

**Authors:** Anthony T. Bogetti, Jeremy M. G. Leung, John D. Russo, She Zhang, Jeff P. Thompson, Ali S. Saglam, Dhiman Ray, Rhea C. Abraham, James R. Faeder, Ioan Andricioaei, Joshua L. Adelman, Matthew C. Zwier, David N. LeBard, Daniel M. Zuckerman, Lillian T. Chong

## Abstract

We present six advanced tutorials instructing users in the best practices of using key new features and plugins/extensions of the WESTPA 2.0 software package, which consists of major upgrades for enabling applications of the weighted ensemble (WE) path sampling strategy to even larger systems and/or slower processes. The tutorials demonstrate the use of the following key features: (i) a generalized resampler module for the creation of “binless” schemes, (ii) a minimal adaptive binning scheme for more efficient surmounting of free energy barriers, (iii) streamlined handling of large simulation datasets using an HDF5 framework, (iv) two different schemes for more efficient rate-constant estimation, (v) a Python API for simplified analysis of WE simulations, and (vi) plugins/extensions for Markovian Weighted Ensemble Milestoning and WE rule-based modeling at the system biology level. Applications of the tutorials range from atomistic to residue-level to non-spatial models, and include complex processes such as protein folding and the membrane permeability of a drug-like molecule. Users are expected to already have significant experience with running conventional molecular dynamics simulations and completed the previous suite of WESTPA tutorials.

## 1 Introduction and Scope of Tutorials

The WESTPA (Weighted Ensemble Simulation Toolkit with Parallelization and Analysis) software package is a highly scalable implementation of the weighted ensemble (WE) path sampling strategy [1, 2] that has helped transform what is feasible for molecular simulations in the generation of pathways for long-timescale processes (> μs) with rigorous kinetics. Among these simulations are atomically detailed simulations of protein folding [3], protein-protein binding [4], protein-ligand unbinding [5], and the large-scale opening of the SARS-CoV-2 spike protein [6]. The latter involved the slowest process (seconds-timescale) yet studied for a massive system (one million atoms) using WE simulations. As a “bleeding edge” application, these efforts have motivated major upgrades to WESTPA (version 2.0) that enable the sampling of processes at even longer timescales and more streamlined handling of large datasets [7]. Like its predecessor, WESTPA 2.0 is a Python package that is (i) interoperable, enabling the use of any type of stochastic dynamics simulation (e.g., MD or Monte Carlo simulations) and any model resolution (e.g., atomistic, coarse-grained, non-spatial or spatially resolved systems biology models) [8, 9]; and (ii) extensible, making it straightforward to modify existing modules or create plug-ins in order to support new scientific efforts.

Here, we present a suite of six advanced tutorials for applying major upgrades in the WESTPA 2.0 software package. For general prerequisites to attempting these tutorials, please see **Section 1.2** below. Among these tutorials is one involving the Markovian Weighted Ensemble Milestoning (M-WEM) approach [10], which interfaces the WE strategy with another path sampling method called milestoning [11, 12]. In the final tutorial, we broaden the scope of path sampling to a systems biology application involving a WESTPA plugin for enhancing the efficiency of Monte Carlo simulations using the BioNetGen systems biology package [13, 14]. All files for the tutorials can be found online in the WESTPA 2.0 GitHub repository https://github.com/westpa/westpa2_tutorials. In each tutorial, we outline learning objectives and expected outcomes.

After completing **Advanced Tutorial 3.1**, which involves the simulation of Na^+^/Cl^−^ association, the user should be able to:

1. Create a customized binless resampler scheme for splitting and merging trajectories based on by k-means clustering using the `BinlessMapper` resampler module;
2. Initiate a WE simulation from multiple starting conformations;
3. Combine multiple WE simulations for analysis using the `w_multi_west` multitool;
4. Perform post-simulation analysis using the `w_crawl` tool.

After completing **Advanced Tutorial 3.2** involving the simulation of drug membrane permeation, the user should be able to:

1. Set up a double membrane bilayer system for permeability studies;
2. Use the highly scalable HDF5 framework for more efficient restarting, storage, and analysis of simulations;
3. Apply the minimal adaptive binning (MAB) scheme.

After completing **Advanced Tutorial 3.3** involving the simulation of ms-timescale protein folding, the user should be able to:

1. Apply the haMSM plugin for periodic reweighting of simulations;
2. Use the `msm_we` package to build an haMSM from WE data;
3. Estimate the distribution of first passage times.

After completing **Advanced Tutorial 3.4** involving the creation of custom analysis routines and calculation of rate constants, the user should be able to:

1. Access simulation data in a `west.h5` file using the high-level `Run` interface of the `westpa.analysis` Python API and retrieve trajectory data using the `BasicMDTrajectory` and `HDF5MDTrajectory` readers;
2. Access steady-state populations and fluxes from the `assign.h5` and `direct.h5` data files, convert fluxes to rate constants, and plot the rate constants using an appropriate averaging scheme;
3. Apply the RED analysis scheme to estimate rate constants from shorter trajectories;

After completing **Advanced Tutorial 3.5** involving simulations of alanine dipeptide using the M-WEM method, the user should be able to:

1. Install the M-WEM software and perform a M-WEM simulation;
2. Create milestones to define the M-WEM progress coordinate;
3. Analyze an M-WEM simulation to compute the mean first passage time, committor, and free energy landscape.

After completing **Advanced Tutorial 3.6** involving rule-based modeling of a gene switch motif using the WESTPA/BNG plugin, the user should be able to:

1. Install the WESTPA/BNG plugin and set up a WESTPA/BNG simulation;
2. Apply adaptive Voronoi binning, which can be used for both non-spatial and molecular systems;
3. Run basic analyses tailored for high-dimensional WESTPA simulations.

In each tutorial, all of the required software, including the dynamics engine and analysis tools, are freely available with easily-accessible online documentation. Please note the version of each software package listed in the **Computational Requirements** section of each tutorial.

### 1.1 The Weighted Ensemble Strategy

WE is a highly-parallel path sampling strategy for generating rare events, for studying non-equilibrium steady states, and less commonly, for studying equilibrium properties. At heart, it is a simple and flexible strategy which is agnostic to system type and which therefore lends itself to numerous applications and optimizations. The properties of WE, including strengths and limitations, have been reviewed in detail before [2], although improvements continue to be developed [16–20]. Here, we briefly review key aspects of WE.

#### The Basic WE Procedure

See **Figure 1**. WE orchestrates multiple trajectories—each assigned a weight—run in parallel by stopping them at regular time intervals of length *τ* (typically a large multiple of the underlying simulation time step), examining the trajectories, and restarting a new set of trajectories. The new trajectories are always continuations of the existing set, but some trajectories may not be continued (they are “pruned”) and others may be replicated. Discontinued trajectories result from probabilistic “merge” events where a continued trajectory absorbs the weight of one that is pruned. Replicated trajectories are said to be “split” with the original weight shared equally among the copies. Usually bins in configuration space are used to guide split and merge events based on a target number of trajectories for each bin, but any protocol—including binless strategies highlighted below—may be used for this purpose. Regardless of the resampling protocol, a “recycling” protocol often is used whereby events reaching a user-specified target state are reinitiated according to a specified distribution of start states [21]. This recycling protocol focuses all sampling on a single direction of a process of interest and has valuable properties as noted below.

**Figure 1.**
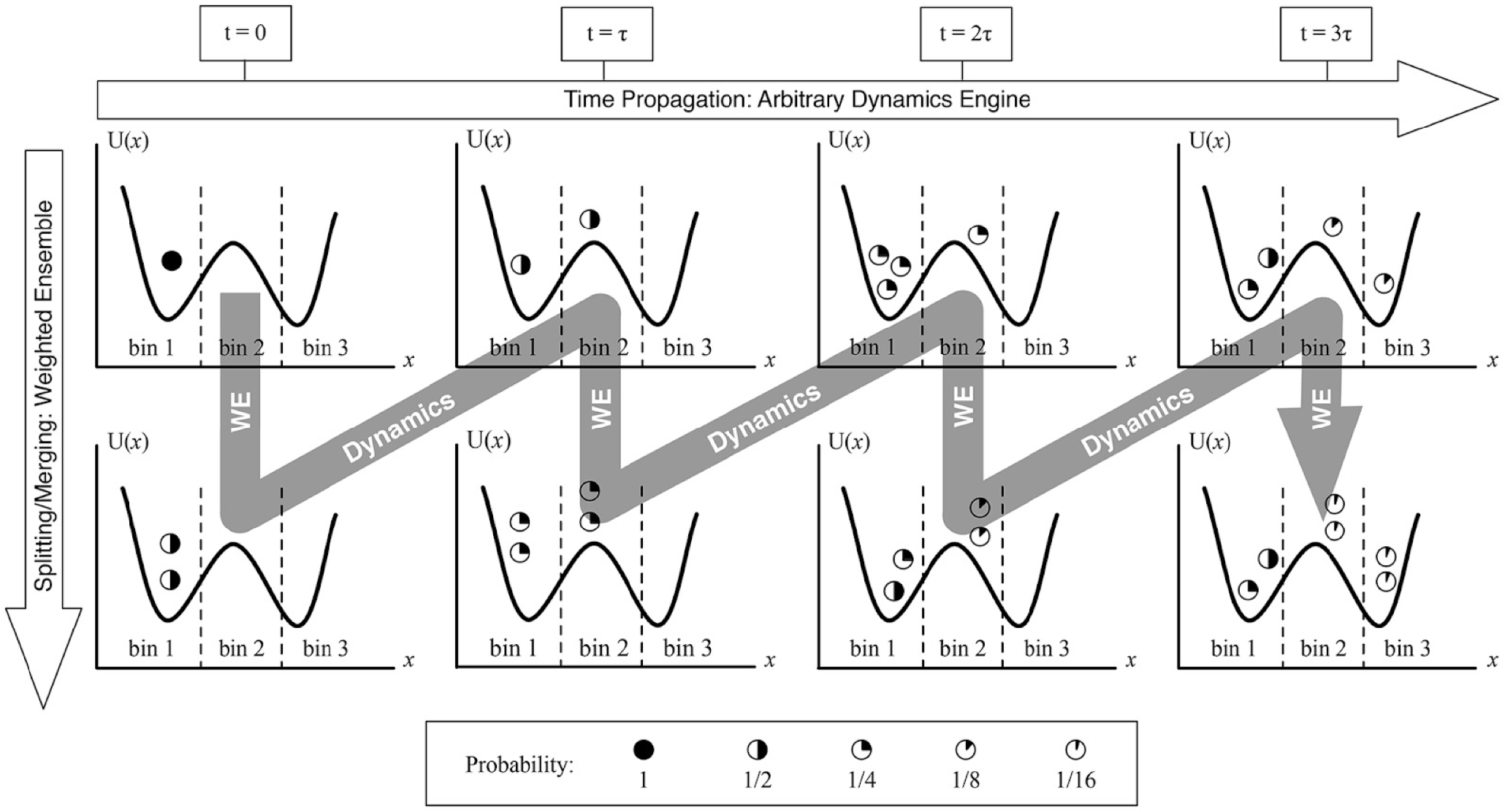
Overview of the weighted ensemble (WE) strategy [9]. WE typically employs bins, demarcated here by dashed vertical lines, to guide a set of trajectories to sample throughout configuration space. Using only ordinary dynamics—without biasing forces—WE replicates (“splits”) trajectories in unoccupied or under-occupied regions of space and prunes (“merges”) trajectories in over-occupied regions, according to the user-specified allocation scheme which here is a target of two trajectories per bin. Throughout the process, weights (partially filled circles) are tracked by statistical rules of inheritance that ensure that the overall ensemble dynamics are consistent with non-equilibrium statistical mechanics [15]. Figure adapted with permission from [8].

#### WE is Resampling, and Hence Unbiased

The simple steps defining WE simulations stem from its basis as a statistical “resampling” procedure [15]. The split/merge steps generate a statistically equivalent (re)sample of an initial trajectory set by increasing/reducing trajectory density in some regions of configuration space at a given time, using weight adjustments to maintain the underlying trajectory distribution. The trajectory set is therefore unbiased at all times, i.e., average time-dependent observables derived from many WE runs will match the average of a large number of conventional simulations without splitting or merging events [15].

Furthermore, the distributions of transition path times (“barrier crossing times”) from WE runs match those from converged conventional simulations, and can be generated in orders of magnitude less computing time [22, 23]. The lack of bias in the dynamics of WE runs holds true regardless of whether recycling is employed.

#### Observables and Ensembles Sampled by WE

WE can yield transient and/or steady-state observables. When recycling is not used, WE provides pathways, i.e., sequences of conformations in a transition and the frequencies of those sequences, in addition to time-dependent observables as the system relaxes to equilibrium, e.g., the probability of a given event at a given time after initiation in the chosen starting state. Complex systems are unlikely to relax fully to equilibrium during a WE simulation. With a recycling protocol, the system will not relax to equilibrium but instead to a non-equilibrium steady state (NESS) that has steady probability flow from initial to target state. If reached, the NESS provides a simple mechanism for computing rate constants via the Hill relation [21]. However, although relaxation to a NESS can be considerably faster than relaxation to equilibrium [24, 25], the process may be too slow for WE to reach NESS on practical timescales, motivating the haMSM approach [3, 17] described below.

#### Resampling Introduces Correlations, which Increase Variance

WE has intrinsic limitations, like any method [26], and it is essential to understand them. Most fundamentally, splitting and merging introduce correlations into the sampled trajectory ensemble that could decrease its information content. These stem primarily from splitting events: multiple trajectories share an identical history up to the time of the split event and hence do not contribute fully independent information to any observable. These correlations, in turn, can lead to large run-to-run variance [3] because the trajectory ensemble in each WE run results from a relatively small number of “parent” trajectories which have been split repeatedly. This variance is addressed to some extent by the iterative haMSM protocol described below, and more directly by ongoing mathematical optimizations noted below. Importantly, correlations within WE ensembles lead to significant challenges in quantifying uncertainty [2, 27].

#### Ongiong Efforts at Optimization and Variance Reduction

Because WE is unbiased so long as a correct resampling protocol is used [15], there is an opportunity to reduce the run-to-run variance noted above by improved resampling procedures. In the context of binned WE simulations, both the construction of bins and the number of trajectories per bin can be optimized based on a recently developed mathematical formulation [20, 28] or based on heuristics [18]. Bins do not need to be kept static over time [15, 18, 29]. Optimization approaches are actively being studied and incorporated into WESTPA as appropriate.

#### WE Cannot Solve Every Problem

Despite its great strengths and highly notable achievements [3, 5, 6], users should not assume WE can tackle any problem. Independent of the correlation/variance issues noted above, certain systems will remain too complex for WE given current hardware and algorithms. In every system, there is a minimum transition path time t_TP_ (also called t_b_) [30, 31] for physically realistic events which sets an absolute requirement on sampling required: in a WE run, a set of trajectories exceeding the minimum t_TP_ must be generated, which may be a prohibitive cost. Additionally, even if the necessary computing resources are available, current binning and resampling strategies might not be sufficient to generate events of interest. And finally, even if events of interest are generated, the sampled trajectories may be insufficient for producing observables of interest such as a reliable estimate of the rate constant.

### 1.2 Prerequisites

#### Background Knowledge and Experience

All tutorials here are at the advanced level. Thus, a prerequisite for these tutorials is completion of the Basic and Intermediate WESTPA tutorials [32], which have been updated for use with WESTPA version 2.0 (https://github.com/westpa/westpa_tutorials). Users should already have extensive experience running conventional simulations using the underlying dynamics engine of interest (Amber [33], OpenMM [34], BioNetGen [13], etc.). Prior to applying the WE strategy to their own systems, we suggest that users run multiple conventional simulations to (i) ensure that the preparation of the system and propagation of dynamics is according to best practices (e.g., see [35]), (ii) identify potential progress coordinates and initially define the target state, and (iii) estimate the ns/day on a single CPU/GPU for your system and storage needs for the full-scale WE simulation. We highly recommend that new WESTPA users read this review article [2] and this introduction to non-equilibrium physics of trajectories [36].

#### Software Requirements

The WESTPA 2.0 software is a standard Python package that can be used on any Unix operating system. The software requires Python versions *≥*3.7 and a number of standard Python scientific computing packages. We recommend installing WESTPA either as a PyPI or conda package using miniconda. Both packages provide all required software dependencies and can be installed using one-line commands: (1) `python -m pip install westpa` or (2) `conda install -c conda-forge westpa`. Note that it is a best practice to install WESTPA into an isolated virtual or conda environment, along with the dependencies specific to your project. Due to the use of the MDTraj Python library with the WESTPA 2.0 HDF5 framework, certain modifications to the installation procedure are required for running WESTPA 2.0 on ppc64Ie architectures (e.g., TACC Longhorn or ORNL Summit supercomputers; see https://github.com/westpa/westpa/wiki/Alternate-Installation-Instructions).

WESTPA 2.0 is designed to be interoperable with any dynamics engine, requiring an external dynamics engine to propagate the dynamics in a WE simulation. Please see the prerequisite sections of each tutorial for additional software requirements that are specific to that tutorial.

#### Hardware Requirements

Like its predecessor, WESTPA 2.0 is highly-scalable on CPUs/GPUs, making optimal use of high-performance computing (HPC) clusters available at academic institutions or supercomputing centers. Memory requirements are dependent on the underlying dynamics engine, e.g., ~1 GB per CPU core (or GPU) for atomistic MD simulations. Users should refer to the best practices of their dynamics engine of choice to determine the optimal allocation of resources for each CPU/GPU. The most efficient way to run WESTPA is to use a computing resource that provides the user with a number of CPUs/GPUs—all the same processor speed—that either matches the number of trajectories per WE iteration or a number by which the number of trajectories at any point in time is evenly divisible. WE can nevertheless run on heterogeneous hardware (different processor or memory bus speeds) or with trajectory counts that do not divide evenly onto CPUs/GPUs, but this scenario decreases efficiency as some processors are inevitably idle for at least a portion of the overall runtime.

Users can estimate the approximate storage space required for their project by taking the product of the following: (i) amount of disk space required for storing data from one trajectory segment of length *τ*, (ii) the maximum number of trajectories per WE iteration, and (iii) the total number of WE iterations required to generate a reasonable maximum trajectory length. To optimize the use of storage space, we recommend that users tar up trajectory files into a single file for each WE iteration and remove coordinates of the system that are not of primary interest (e.g., solvent coordinates for certain processes). We note that the WESTPA 2.0 HDF5 framework dramatically reduces the storage space required for trajectory coordinates by consolidating the data from millions of small trajectory files into a relatively small number of larger HDF5 files, reducing the large overhead from the file system that results from the storage of numerous small trajectory files. By doing so, the HDF5 framework also alleviates potential I/O bottlenecks when a large amount of simulation files are written after each WE iteration.

## 2 Recommended Simulation Workflow with WESTPA 2.0 Upgrades

Given the major upgrades in the WESTPA 2.0 software package [7], we recommend the three-stage simulation workflow illustrated in **Figure 2**. Details of the particular schemes are provided in the tutorials.

**Figure 2.**
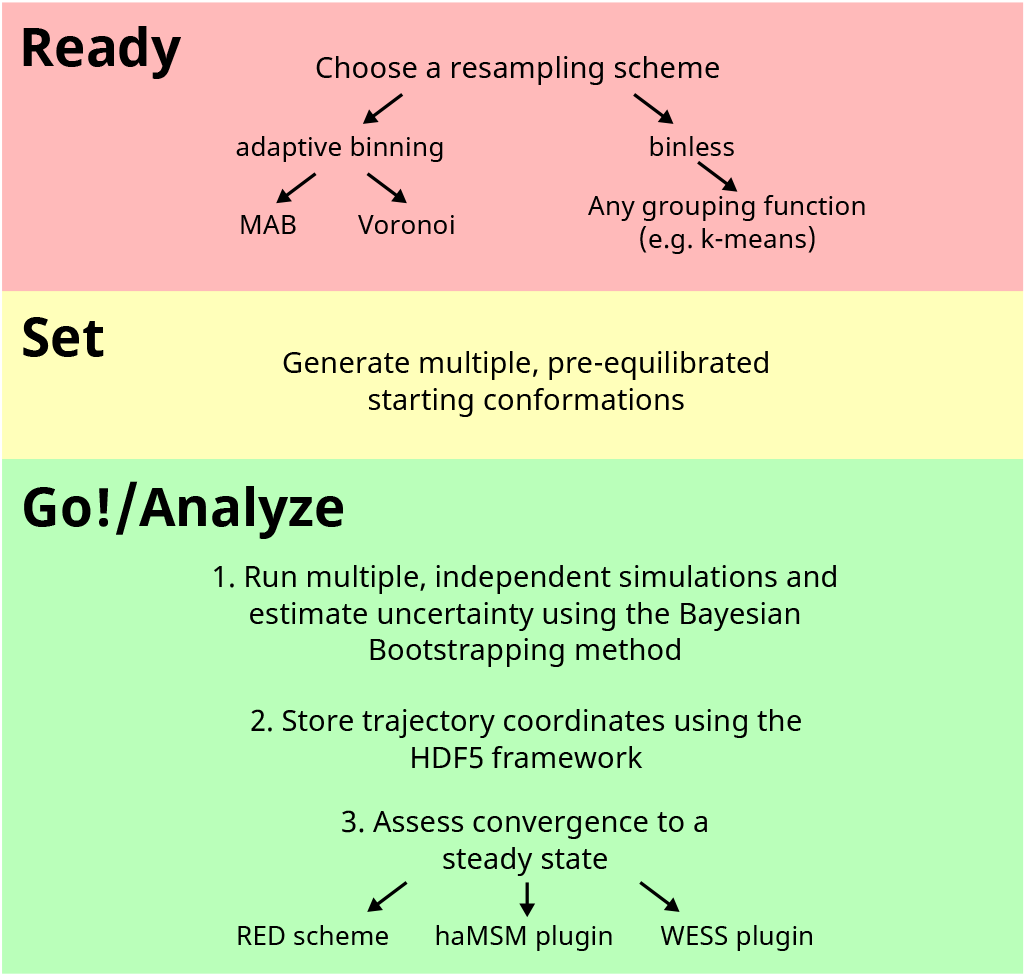
Recommended simulation workflow that makes use of major upgrades in the WESTPA 2.0 software.

**Figure 3.**
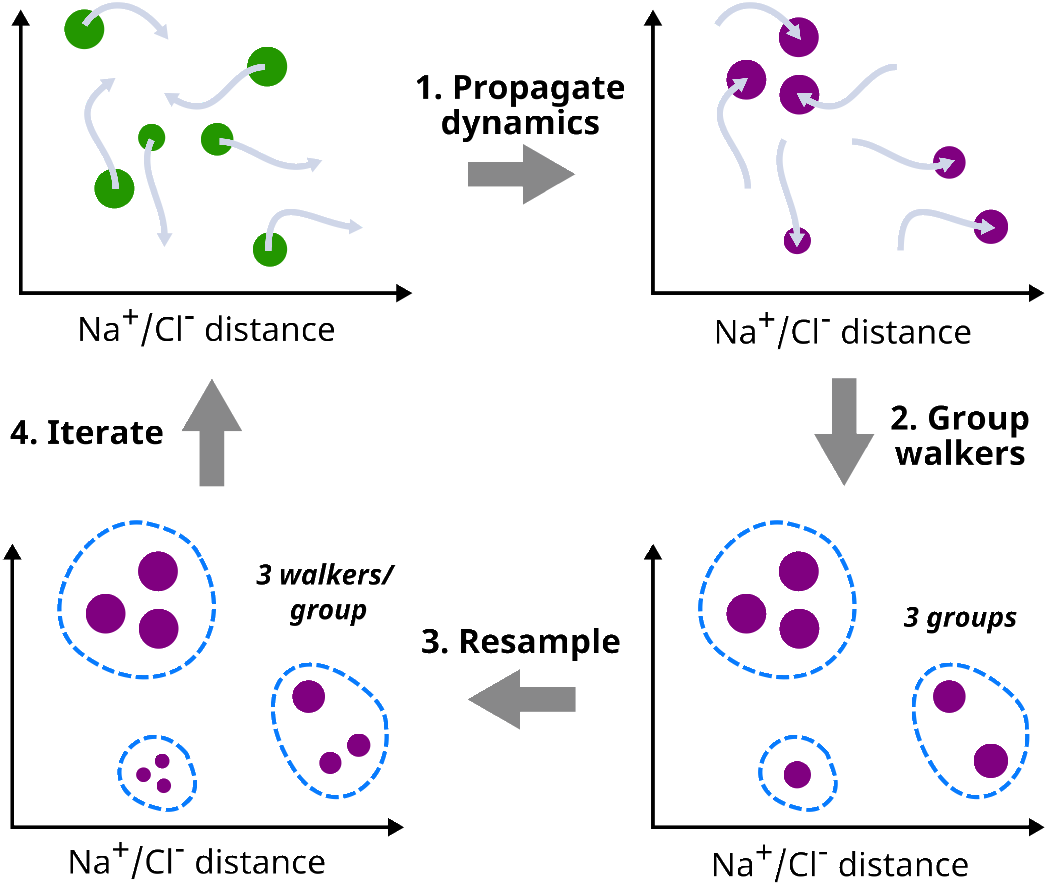
Flow chart for simulating Na+/Classociation using the new binless resampler module. After dynamics propagation in step 1, trajectories in each bin are grouped with the user-specified group.py function. In this example, trajectory walkers are grouped using k-means clustering.

In the first (“Ready”) stage, if one uses a binned resampling scheme, we recommend using one of the two adaptive binning schemes available in WESTPA 2.0: the minimal adaptive binning (MAB) scheme or adaptive Voronoi binning scheme. These adaptive binning schemes enable quicker explorations of the chosen progress coordinate than manual, fixed binning schemes. The MAB scheme is effective at surmounting barriers in a direction of interest [18] while the adaptive Voronoi binning scheme [15] is ideal for enhanced sampling in high-dimensional space (more than three dimensions) when all parts of the progress-coordinate space are potentially important. However, if progress-coordinate space includes, for example, undesirable unfolded protein conformations, adaptive Voronoi binning might allocate bins and computing resources to those regions. Besides the adaptive binning schemes, one can opt for a “binless” resampling scheme by defining a grouping function as described in **Advanced Tutorial 3.1** below.

In the second (“Set”) stage, we recommend starting the WE simulation from multiple, pre-equilibrated starting conformations that are representative of the initial stable state (at least one “basis state” for each trajectory walker) to improve the sampling of the initial state and diversity of generated pathways to the target state.

In the third (“Go/Analyze”) stage, we recommend applying one of the following three options to further accelerate convergence to a steady state once successful pathways are generated. The Rates from Event Durations (RED) analysis scheme [19] estimates rate constants more efficiently than the original WE scheme [1] by exploiting information in the transient region of the simulation. Another option is the haMSM plugin, which employs a fine-grained “microbin” analysis and can be used to not only estimate rate constants following WESTPA simulation (e.g., for the seconds-timescale coronavirus spike opening process [37]), but to also restart trajectories with their weights adjusted for a steady state [7]. Because restarts in the haMSM plugin are initiated from configurations occurring throughout previously run trajectories, the continuity of the generated pathways is broken. The third option is the weighted ensemble steady state (WESS) plugin [21], which uses the less fine-grained WE bins to estimate steady state but preserves the continuity of pathways, restarting from only the final points of trajectories. While all trajectory files of the chosen dynamics engine are saved by default, we recommend storing the trajectory coordinates using the WESTPA 2.0 HDF5 framework, which greatly facilitates the restarting, storage efficiency, and analysis of WE simulations. When possible, users should run multiple WE simulations, which provides a greater number of independent pathways and enables straightforward estimation of error using the Bayesian bootstrap method [27] (see https://github.com/ZuckermanLab/BayesianBootstrap).

### Random Seeds for Simulations

For WE trajectories to diverge from one another after a splitting event, a stochastic thermostat is required for MD simulations. Furthermore, the random number seeds for such thermostats must be sufficiently random (uncorrelated) to avoid undesired bias of the dynamics when trajectories are restarted at short time intervals (e.g., in the case of WE simulations) [38]. To avoid such bias, we strongly recommend using WESTPA’s system entropy-seeded random number facility instead of any time-seeded random number generator of the chosen dynamics engine. To use this facility, we first set the random seed to `RAND` in the dynamics input file (e.g., `ig=RAND` in the AMBER `md.in` file) and then specify this input file in `runseg.sh`, which will replace the `RAND` string with the WESTPA random number seed.

### Extremely Low Trajectory Weights

While it is possible to set a minimum threshold weight (e.g., 10^−100^) for trajectories to be considered for splitting, the generation of trajectories with extremely low weights (e.g., <10^−100^) is a potential warning sign that the division of configurational space is not capturing all relevant free energy barriers. If a WE simulation yields such trajectories, we strongly suggest re-evaluating the choice of progress coordinate and/or restricting the binning to a carefully chosen subset of configurational space that would avoid generating such trajectories. For example of the latter, see **Advanced Tutorial 3.2**.

## 3 Advanced Tutorials

### 3.1 Creating “Binless” Resampling Schemes: Na^+^/Cl^−^ Association Simulations

#### 3.1.1 Introduction

The non-linearity of certain progress coordinates (e.g., those identified by machine learning tools) requires the creation of “binless” rather than binned resampling schemes for rare-event sampling. In addition, binless schemes can be useful in grouping trajectories by a feature of interest for resampling. For example, trajectories could be grouped by history (sharing the same parent structure) to improve the diversity of trajectories that successfully reach the target state, or by a simple k-means clustering. This tutorial builds upon the Basic WESTPA Tutorial (Na^+^/Cl^−^ association simulations) [32] by introducing users to running and analyzing a WESTPA simulation that employs “binless” resampling schemes. This tutorial is a prerequisite for **Advanced Tutorials 3.2-3.4**.

##### Learning Objectives

This tutorial introduces users to the generalized resampler module in the WESTPA 2.0 software package that allows for the creation of either binned or binless resampling schemes.

The tutorial also instructs users on how to initiate a WE simulation from multiple representative conformations of the starting state and how to apply two key post-simulation analysis tools. Specific learning objectives are:

1. How to create a binless scheme for splitting and merging trajectories based on k-means clustering using the `BinlessMapper` resampler module;
2. How to initiate a WE simulation from multiple starting conformations;
3. How to combine multiple WE simulations for analysis using the `w_multi_west` tool;
4. How to perform post-simulation analysis using the `w_crawl` tool.

#### 3.1.2 Prerequisites

##### Computational Requirements

This simulation can be completed in 5 hrs using 8 Intel Xeon W3550 3.07 GHz CPU cores, generating 1 GB of data using the HDF5 framework of the WESTPA 2.0 software package. Two independent `west.h5` files, each containing 100 WE iterations of simulation data, are provided for the analysis portions of this tutorial in the `for_analysis/` directory. This tutorial uses the OpenMM 7.6 software package for dynamics propagation (https://openmm.org) and the MDTraj 1.9.5 analysis suite (https://www.mdtraj.org/1.9.5/index.html) for calculations of the WE progress coordinate. The scikit-learn 1.1.0 package (https://scikit-learn.org) is used to identify the “binless” groups.

##### Jupyter Notebook

Sample code for running and analyzing a WESTPA simulation according to “best practices” is made available in a Jupyter notebook. For the visualization portions of the notebook, nglview, matplotlib, and ipympl are required.

##### Quick Start for this Tutorial

Users can run the following command within the `tutorial-3.1/` directory to install all the software dependencies for this tutorial to an existing conda environment:

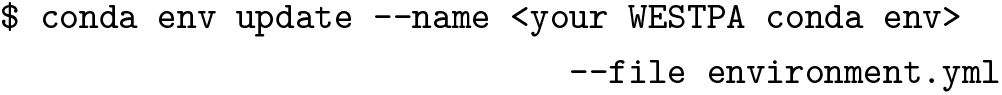

#### 3.1.3 Setting up the WE Simulation

This simulation uses the same WE parameters (*τ*, number of trajectories per bin, etc.) as the Basic WESTPA Tutorial (Na^+^/Cl^−^ association simulations) [32]. The following are major differences from the Basic Tutorial that highlight more advanced features of WESTPA simulation setup.

##### Binless Resampler Module

The binless resampler module can be accessed from the `west.cfg` file in the `system_options` section where binned and binless schemes are all defined. If the `BinlessMapper` is used by itself, the entirety of configurational space will be binless. To recycle trajectories while using a binless framework, we will need to place a `BinlessMapper` inside of a `RecursiveBinMapper` bin and demarcate the target state in a separate RecursiveBinMapper bin. This framework is identical to the one used for the MAB scheme with recycling.

The `BinlessMapper` takes three mandatory arguments. The first, `ngroups`, specifies the total number of groups to assign trajectory walkers in the binless space. The second, `ndims`, specifies the dimensionality of the progress coordinate and is limited to either 1 or 2 at this time. The `ndims` parameter specifies the dimensionality of clustering, e.g., 2 for generating clusters in two-dimensional space (each dimension will not be grouped separately as is typically the case for binned resampling schemes). The clustering of trajectories enables sampling of high-dimensional space without an exponential increase in the number of walkers. The final argument is `group_function`, which specifies the function for grouping trajectories in an external file (here, `group.py`) and will take as input the progress coordinates and the `ngroups` values. We provide a general example of using this function for a one-dimensional progress coordinate using k-means clustering. Additional keyword options can be specified under `group_arguments` in the `west.cfg` file. An example of using a recursive binless scheme is shown in the `west.cfg` snippet below:

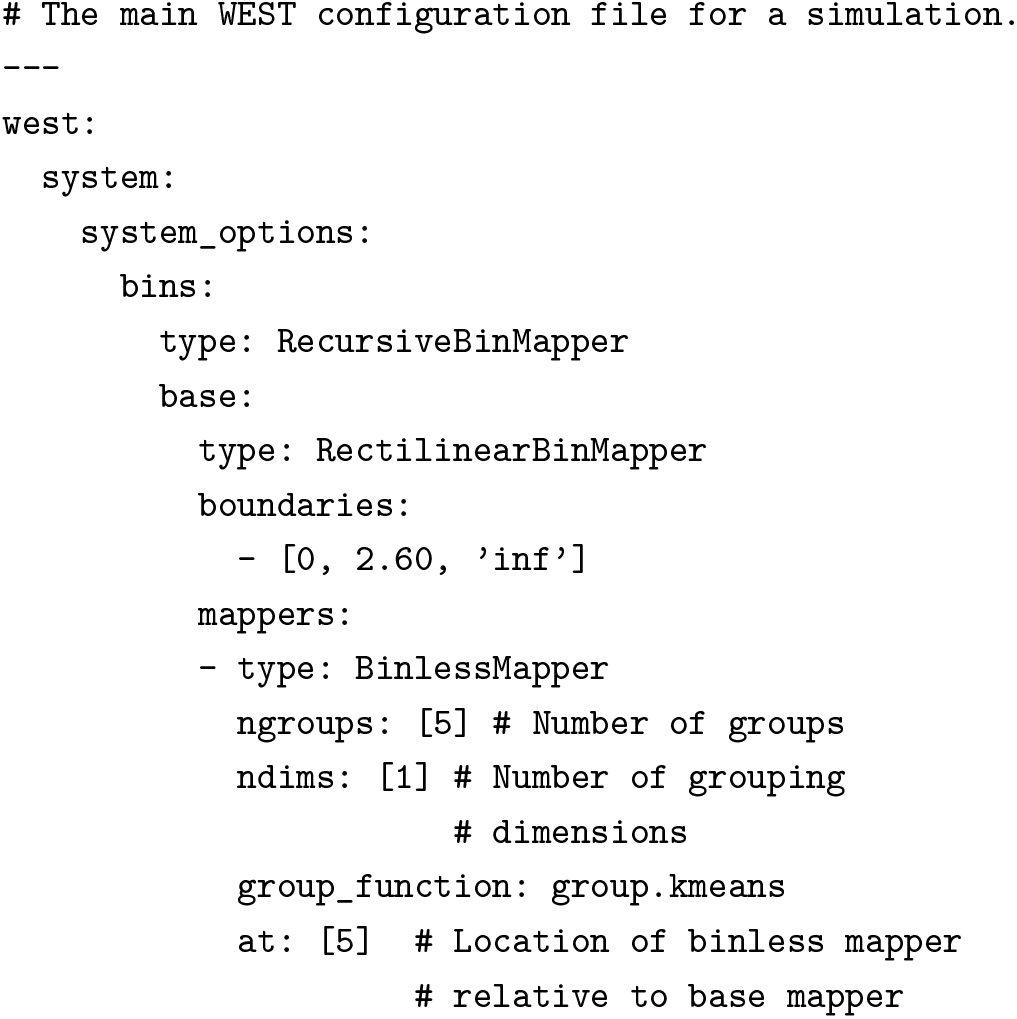

##### Initiating a WE Simulation from Multiple Structures

Ideally, a WE simulation is initiated from multiple preequilibrated structures that are representative of the initial state for the rare-event process of interest, e.g. using a conventional simulation or a separate WE simulation of the initial state. Within the WESTPA framework, we refer to these structures as “basis states”. If the simulation is run under non-equilibrium steady-state conditions, trajectories that reach the target state are “recycled” by terminating that trajectory and initiating a new trajectory with the same statistical weight from one of the basis states. Structural files (xml files in this tutorial) for basis states contain the coordinates and velocities, and are placed in separate, numbered folders within the `bstates/` directory. Accompanying each xml file is a `pcoord.init` text file which contains the progress coordinate value of that basis state. These progress coordinates are saved to the HDF5 file during the initialization process. The `bstates/` directory also contains a reference file (`bstates.txt`) that lists all of the available basis states. The `bstates.txt` file is formatted with three columns, corresponding to the basis states’ names, associated probabilities, and folder name, respectively. Additional basis states can be added as separate, additional lines at the end of the `bstates.txt` file. The probability over all basis states must sum to one, and will be normalized by WESTPA during the initialization process to sum to one if the condition is not met. Compared to the Basic WESTPA tutorial involving Na^+^/Cl^−^ association [32], the `get_pcoord.sh` and `runseg.sh` files are also modified such that `$WEST_DATA_STRUCT_REF` now corresponds to the directory for each basis state and not the xml file itself.

#### 3.1.4 Running the WE Simulation

As with previous versions of WESTPA, the simulation can be initialized using `./init.sh` and run using `./run.sh`. Alternatively, both steps could be executed consecutively using the new Python API by running the following command:

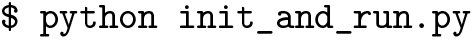

The `init_and_run.py` script will print out simulation updates to the console in real-time. An example `runwe.slurm` file with commands for both methods of execution is provided for use with SLURM-like workload managers. We also provide a Jupyter notebook that demonstrates the steps for cleaning up, initializing, and running the WE simulation.

Note that this tutorial is using the new HDF5 trajectory storage framework, which will be explained further in **Advanced Tutorial 3.2**. To enable the use of the HDF5 framework in your own simulation, you may use the current tutorials directory as an example. The location of the trajectory h5 files will need to be specified in the `west.cfg` file, and the appropriate restart and topology files will need to be copied to the locations specified in the `get_pcoord.sh` and `runseg.sh` files. You will also need to make sure that the file extensions for any trajectory files are readable by `mdtraj.load`, (e.g., Amber restart files must end in .ncrst) which simply requires renaming. To save disk space, trajectory files outputted by the dynamics engine can be deleted after every iteration in the `post_iter.sh` file, which is located in the `westpa_scripts/` directory.

#### 3.1.5 Monitoring and Analyzing the WE Simulation

##### Combining Multiple WE Simulations for Analysis

To combine multiple WE simulations into a single aggregate simulation file for analysis, we can use the `w_multi_west` tool, which creates a single `multi.h5` file that contains the data from all of the `west.h5` files of each WE simulation. Each WE iteration in the `multi.h5` file contains all of the trajectory segments from the corresponding iterations of the individual WE simulations, all normalized to the total weight for that iteration. For backwards compatibility, a version of `w_multi_west` for use with previously run WESTPA 1.0 simulations (v2020.XX) has been available since version 2020.04.

To apply the multitool to a combination of `west.h5` files, place the `west.h5` file for each simulation in a numbered directory starting with `01/`. If all of the simulations used a custom grouping function (such as in `group.py`), you must also include that file in the top-level analysis directory.

The files will be organized as follows:

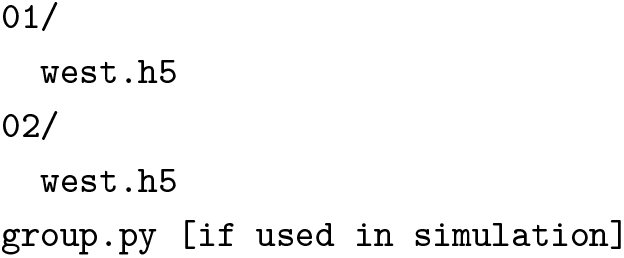

Next, in this directory, run the following to merge the `west.h5` files:

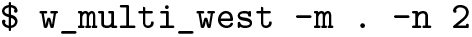

The `-m` flag specifies the path to your directories and the `-n` flag specifies the number of WE simulations to combine for analysis. To combine auxiliary datasets, one can add either an `--aux=NAME_OF_DATASET` flag for a specific dataset or an `--auxall` flag for all auxiliary datasets; note that the inclusion of auxiliary datasets will substantially extend the time needed to combine the simulation data. The above `w_multi_west --auxall` command will generate a list of WE simulation datasets to combine based on the datasets listed in `01/west.h5` and generate a `multi.h5` file with the combined simulation datasets. You may want to rename this file to `west.h5` in order to apply the `w_pdist` tool to the combined simulation dataset. Note that the `w_multi_west` tool will only merge up to the *N-1* WE iteration, ignoring the last WE iteration. The resulting `multi.h5` file will not link to the individual iteration HDF5 files generated using the HDF5 framework.

##### Post-Simulation Analysis

As mentioned above, WESTPA 2.0 enables efficient post-simulation analysis of trajectory data by storing trajectory data in highly compressed HDF5 files. The `w_crawl` tool can then be used to “crawl” through the trajectory data in single HDF5 files per WE iteration rather than millions of trajectory files. Results from the analysis are written to a dataset in a new HDF5 file. Before crawling through an entire simulation dataset, we recommend that users first test their analysis scheme in the `wcrawl_functions.py` file to ensure that the scheme works as expected. For this tutorial, we will only be using a single CPU core for these `w_crawl` calculations but also include a sample script as an example of how to use `w_crawl` on multiple CPU cores in parallel. The `wcrawl_functions.py` file contains the main analysis code. This script first identifies the final frame of a segment’s parent trajectory file from the previous WE iteration and makes sure this is eventually combined with the trajectory segment from the current WE iteration. The inclusion of the parent structure at the beginning of the current iteration trajectory is necessary for using the crawled dataset with WESTPA’s kinetics analysis tools. In this example, trajectory coordinates of only Na^+^ and Cl^−^ are extracted using the MDTraj analysis suite and multiplied by 10 to convert from nm to Å. Resulting per-iteration coordinate values are then saved to an array, which is subsequently saved to a `coord.h5` file. The `coord.h5` file is formatted similarly to a `west.h5` file, where the new per-iteration values are stored under `iterations/iter_{n_iter:08d}/coord`. To ease analysis, a `copy_h5_dataset.py` script is provided to copy `coord.h5`’s contents into a west.h5 file as an auxiliary dataset. Note that if you store a WESTPA simulation’s trajectory HDF5 files in a separate directory from what is used in this tutorial, you need to specify the directory where the `iter_XXXXXX.h5` files are located in the `wcrawl_functions.py` file.

The `run_w_crawl.sh` shell script runs the `w_crawl` tool at the command line and provides options for running the tool in serial or parallel modes. In this tutorial example, we will run the `w_crawl` tool in the serial mode using the `--serial` flag, analyzing one WE iteration at a time on a single CPU core. While the serial mode is sufficient for “crawling” relatively small datasets, the parallel mode using the `--parallel` flag is desirable for datasets with over 100 trajectory segments per WE iteration and/or hundreds of WE iterations. In the parallel mode, each CPU core of a single compute node analyzes a different WE iteration at the same time. To run the `w_crawl` tool across multiple nodes, one can use the ZMQ work manager. Once satisfied with the `wcrawl_functions.py` and `run_w_crawl.sh` files, run the `w_crawl` tool locally:

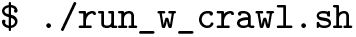

or on a multi-node cluster using the Slurm workload manager:

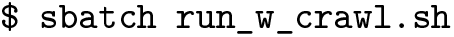

To monitor the progress of the analysis, we examine the `w_crawl.log` file, which contains analysis results for each WE iteration and each trajectory segment. Finally, to copy the `coord.h5` file to the `west.h5` file, run the `copy_h5_dataset.py` script.

#### 3.1.6 Conclusion

After completing this tutorial, users will gain an understanding of how to configure the upgraded resampler module for using a binless scheme, initiate a WE simulation from multiple structures using the new HDF5 trajectory storage framework, and apply the `w_multi_west` and `w_crawl` post-simulation analysis tools.

### 3.2 Simulations of Membrane Permeation by 1-Butanol

#### 3.2.1 Introduction

The ability of a drug-like molecule to cross (or permeate) a lipid bilayer has been of great interest to drug discovery [39], but is challenging to simulate due to the long timescales involved. In this tutorial, we will use WESTPA 2.0 to simulate pathways for membrane permeation by a small molecule (1-butanol) and calculate the permeability coefficient. Our WE protocol employs and explains two new features in WESTPA 2.0 [7]: (i) the Minimal Adaptive Binning (MAB) scheme [18], and (ii) the HDF5 framework for efficient restarting, storage, and analysis of a WE simulation.

##### Learning Objectives

This tutorial demonstrates how steady state WE simulations can be used to generate pathways and permeability coefficients for membrane permeation by a small molecule. Specific learning objectives include:

1. How to set up a double membrane bilayer system for permeability studies;
2. How to use the highly scalable HDF5 framework for more efficient restarting, storage, and analysis of simulations;
3. How to apply the minimal adaptive binning (MAB) scheme.

#### 3.2.2 Prerequisites

In addition to completing the Basic and Intermediate WESTPA Tutorials [32], a prerequisite to this advanced tutorial is completion of the above **Advanced Tutorial 3.1**. Also required is a working knowledge of the CHARMM-GUI membrane builder, PACKMOL, OpenEye Scientific’s OEChem and Omega toolkits (for system preparation only), MDTraj analysis suite, and the OpenMM 7.6 dynamics engine (for running WE).

##### Computational Requirements

The membrane permeability tutorial simulation runs best using, at minimum, a dual-GPU workstation. For this tutorial, simulations were tested with a compute node containing both a NVIDIA Titan X (Pascal) GPU and a NVIDIA GTX 1080 GPU, as well as a 16-core Intel Xeon X5550 CPU running at 2.67 GHz with a total of 100 GB of system memory. In the case a user does not have a GPU and only CPUs, switch between OpenMM’s GPU and CPU platforms by changing the platform name in line 22 of `memb_prod.py` to `CPU` instead of `CUDA`. The complete tutorial simulation run length (37 iterations) required ~4 days of continuous wall clock time on both GPUs, as well as ~30 GB of hard disk space with the HDF5 framework and MAB options turned on.

This tutorial uses the OpenMM 7.6 dynamics engine [34] and MDTraj 1.9.5 analysis suite (https://www.mdtraj.org/1.9.5/index.html) for progress coordinate calculations. Force fields used in this tutorial can be installed via openmmforcefields (https://github.com/openmm/openmmforcefields). System setup and equilibration were performed separately using OpenMM. In order to run the companion Jupyter notebook, nglview, matplotlib are required for visualization purposes. Other dependencies, including NumPy and MDTraj, are installed through WESTPA 2.0 itself.

##### Quick Start for this Tutorial

Users can run the following command within the `tutorial-3.2/` directory to install all the software dependencies to an existing conda environment:

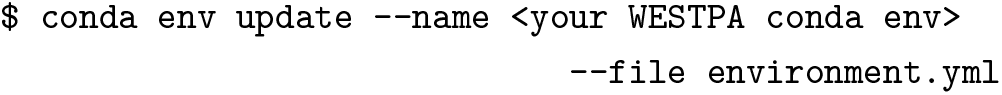

#### 3.2.3 Preparing the simulation

The following preparation steps have already been completed and are presented to instruct the reader on how to prepare similar systems for WE simulation.

Our system consists of a 1-palmitoyl-2-oleoyl-snglycero-3-phosphatidylcholine (POPC) membrane bilayer. The 1-butanol-double POPC membrane bilayer system was prepared by piecing together several smaller molecular systems in the following way. First, a single POPC membrane bilayer was generated using CHARMM-GUI’s membrane builder with 50 lipids per leaflet and zero salt concentration. This membrane was then equilibrated using a single GPU with the OpenMM dynamics engine using the standard CHARMM-GUI procedure. The membrane plus the outer aqueous layer to the membrane, once combined, (see **Figure 4**) was equilibrated for an additional 500 ns. A 2D representation of 1-butanol was generated from an input SMILES string (CCCCO) using OEChem, and converted to a 3D structure using the Omega TK toolkit. The 3D structure of 1-butanol was then solvated with a 2 nm slab of water molecules at a density of 1 gm/cm^3^ using PACKMOL along with the OEChem TK and Omega TK toolkits from OpenEye. Finally, the full double-membrane bilayer system was assembled by placing the butanol-embedded slab of water at the origin, with a single-membrane system at each z-edge of the water slab. The resulting system was then subjected to energy minimization and equilibrated before the initiated WE simulations of butanol permeating the membrane bilayer were initialized.

**Figure 4.**
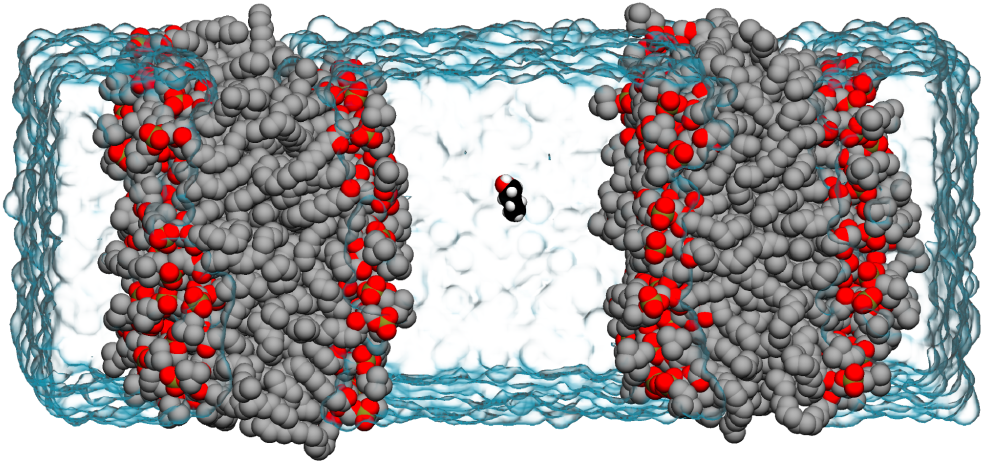
The equilibrated double-membrane bilayer system used for initiating WE simulations of membrane permeation by 1-butanol in this tutorial. Both the POPC membrane bilayers (gray; hydrogens removed for clarity) and 1-butanol (black) are represented in van der Waals representation, while layers of explicit water molecules are shown as a transparent blue surface.

##### The System

In this tutorial, we will run a WE simulation of 1-butanol crossing one membrane of a double POPC membrane bilayer system. To run the WESTPA 2.0 simulation, the AMBER LIPID17 force field is applied to all POPC lipids, explicit water molecules are represented by the TIP3P model, while the parameters for 1-butanol were taken from the GAFF 2.11 force field. All force field parameters were applied using the openmmforcefields Python package.

##### Progress Coordinate

The progress coordinate (z) is defined as the (signed) distance from the center of mass (COM) of the butanol molecule to that of the closest membrane. The width of a single leaflet of the membrane is roughly 2 nm, so if z *≤* 2 nm, between −2 and 2 nm, or >2 nm indicates that the butanol molecule has not crossed, is crossing, or has crossed the membrane. A target state of z *≥*3.5 nm is used to recycle trajectory walkers. The actual computation is performed by measuring the signed distances between the COMs of butanol and each of the two membranes, z_1_ and z_2_, using the MDTraj analysis suite and then taking the larger value of the two, z=max(z_1_, z_2_).

##### Preparing the Simulation Environment

Once we have constructed and equilibrated the 1-butanol membrane system, we will prepare the WESTPA system environment. First, we will analyze the equilibrated double membrane bilayer system to define an initial progress coordinate. The progress coordinate, equilibrated coordinate file (e.g. XML file), and `bstates.txt` file describing the initial basis states are placed in the bstates/ directory. Second, we will edit the `west.cfg` file with options for using the MAB scheme and HDF5 framework. To initialize the WESTPA 2.0 environment, we will run `./init.sh`. This command will source the WESTPA 2.0 environment, construct the `seg_logs/, traj_segs/`, and `istates/` directories, and will run the `w_init` command with the correct settings for the target state (z = 3.5 nm) and the basis states constructed above.

##### Adaptive Binning using the MAB Scheme

By default, this tutorial uses a manual, fixed binning scheme, but can be modified to use the Minimal Adaptive Binning (MAB) module, which adaptively positions bins along the progress coordinate. To enable this adaptive binning scheme, uncomment the MAB-related lines in the `west.cfg` file, which specify the `MABBinMapper` as the primary bin mapper type, and comment the lines related to the inner `RectilinearBinMapper`. Next, define `n_bins` (the number of MAB bins placed per progress coordinate dimension) in the same section of the `west.cfg` (e.g., if a two-dimensional progress coordinate is being used, [20, 20] indicates 20 bins in each dimension). It is important to note that if the recycling of trajectories at a target state is desired within the MAB framework, recursive bins must be specified by adding a `MABBinMapper` inside of a `RecursiveBinMapper` outer bin and defining the target state in terms of the recursive outer bins. An example of a MAB recursive binning scheme is shown below:

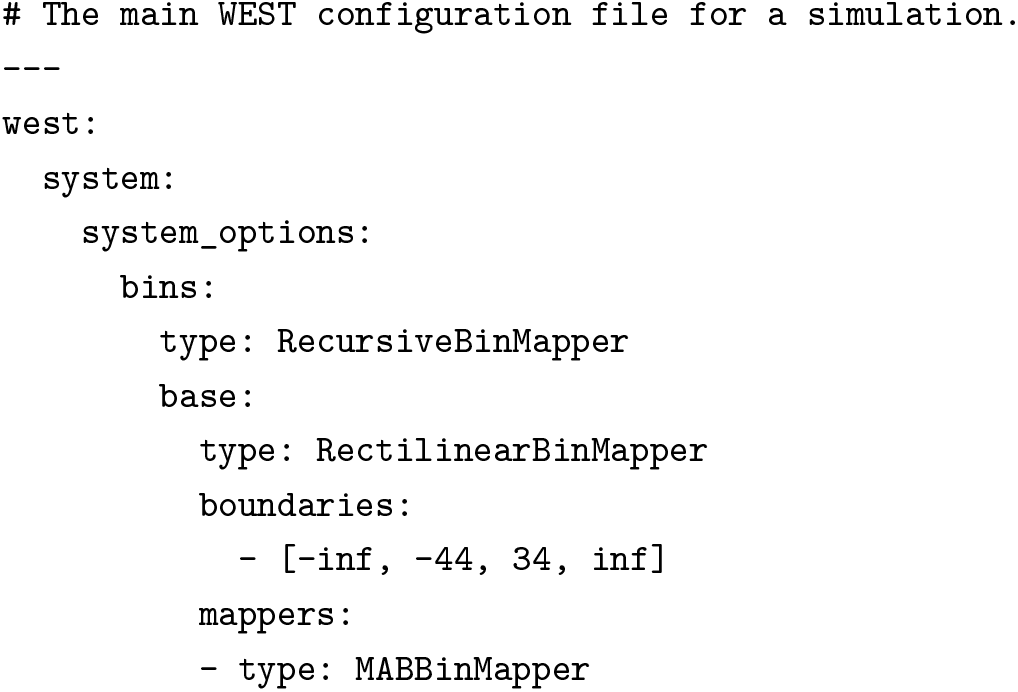
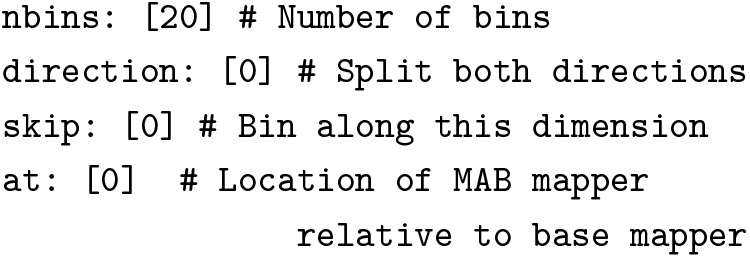

In the above example, the `at` option in the last line specifies which outer bin to place the MAB scheme inside of range `[-44, 34]`. For a two-dimensional progress coordinate, this option will require a list with two values, one for each dimension. The `nbins` option specifies the number of MAB linear bins that will be used inside the bin, plus two more bins for extrema and bottleneck trajectories, respectively. Optional `direction`, `skip` and `mab_log` parameters can also be specified for the MAB scheme. The `direction` parameter (0, −1, or 1) can be used to specify the direction along the progress coordinate for splitting of trajectories, where 0 indicates both directions, −1 indicates the direction of decreasing values along the progress coordinate, and 1 indicates the direction of increasing values along the progress coordinate. The `skip` parameter (1 or 0) designates whether a particular dimension along the progress coordinate will be binned during the simulation, but will be used to define the target state (1 indicates that the dimension will be skipped for binning and 0 indicates that the dimension will not be skipped for binning). The `mab_log` parameter, when enabled with true, will print MAB-related statistics such as the progress coordinate values of extrema walkers to the `west.log` file. Multiple MAB schemes can be added to a recursive binning setup, but only one MAB scheme may be used per each outer bin.

If users choose to combine the application of the MAB scheme with the weighted ensemble steady-state (WESS) plugin [21], which reweights trajectories towards a non-equilibrium steady state, they must provide fixed bins for the reweighting procedure. The positions of these fixed bins can be specified in the WESS plugin section of the `west.cfg` file:

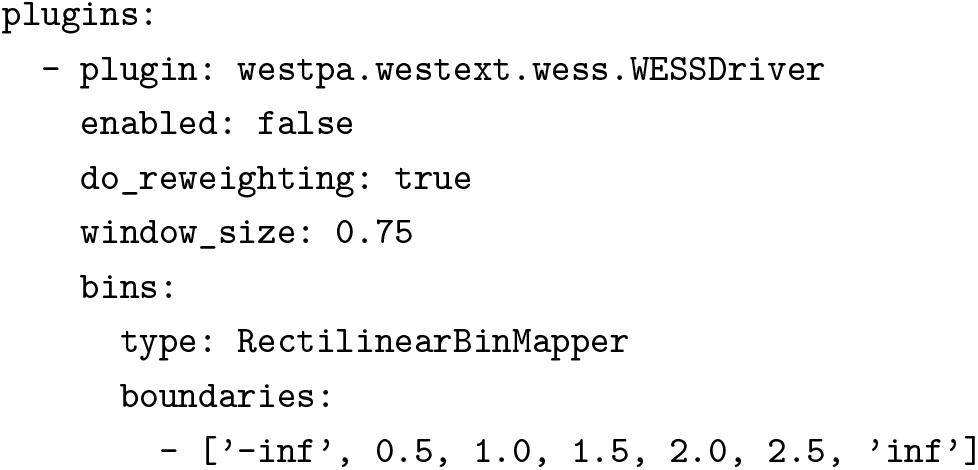

##### HDF5 Framework

The setup for a WESTPA simulation with the HDF5 framework is similar to a vanilla one with the addition of the following procedures, which are a more detailed list of the same steps discussed briefly in **Advanced Tutorial 3.1** above:

1. An iteration entry was provided in `west.cfg` under `west.data.data_refs` to specify where and how the per-iteration HDF5 files should be saved and named.
2. All the necessary files needed for propagating the next segment, such as state/restart and topology files, are passed on to WESTPA through the environment variable, `$WEST_RESTART_RETURN`, after initialization and propagation of each iteration (**Figure 5A**). This information is typically placed in the `get_pcoord.sh` and `runseg.sh` files.
3. All the trajectory files, and topology files if the topology is not stored as part of the trajectory file, are provided to WESTPA through the environment variable, `$WEST_TRAJECTORY_RETURN`, after the propagation of each iteration. This, again, is typically placed in the `get_pcoord.sh` and `runseg.sh` files. The coordinates of the basis states can be provided through the environment variable during initialization to be stored as the “trajectories” of the zero-th iteration. Note that the procedures described in step 2 and 3 are similar to how the progress coordinates are returned through `$WEST_PCOORD_RETURN` in the vanilla WESTPA simulation. The trajectory and restart files will be saved as part of the per-iteration HDF5 files. In turn, these files do not need to be located and copied over to the directory for propagating the next segment, and they will be automatically extracted and put into the segment folder by WESTPA instead (**Figure 5B**).

**Figure 5.**
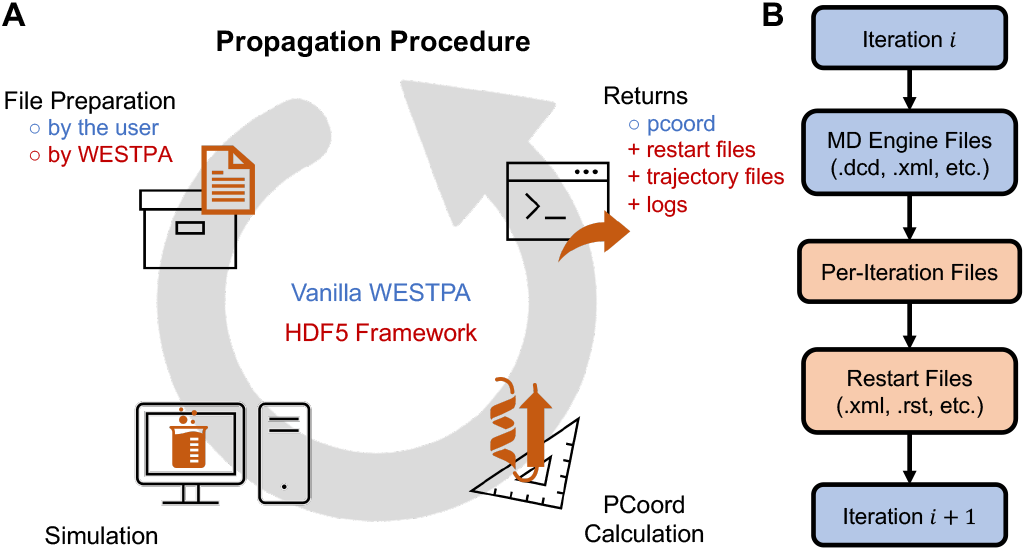
Diagrams showing the differences in the propagation procedure between a vanilla WESTPA run and a run with the HDF5 framework. A) Procedures for propagating one WE iteration using the original WESTPA 1.0 framework (blue text) and WESTPA 2.0 HDF5 framework (red text). Using the WESTPA 2.0 HDF5 framework, the WESTPA software prepares the input files while the user is responsible for returning the progress coordinate (pcoord), restart, trajectory, and optional log files. B) Workflow for using the WESTPA 2.0 HDF5 framework. Blue: Steps and files from the original WESTPA 1.0 procedure. Red: Files generated or prepared by the WESTPA 2.0 HDF5 framework.

These additional procedures simplify the data management on the user’s end for two reasons. First, all the trajectories are stored in a standard way which enables fast and easy access to these trajectories with their associated WE-related data using the newly added analysis module (see Section 3.2.5). Second, the restart/state files are much easier to locate as when they are generated than when they are needed in the next iteration, so letting the users pass trajectory and restart files to WESTPA for it to manage frees users from tracking those files themselves which would require the critical knowledge of how two WESTPA-assigned environment variables work (namely, `$WEST_CURRENT_SEG_INITPOINT_TYPE` and $WEST_PARENT_DATA_REF).

For this membrane permeation example, the per-iteration HDF5 files are saved in a folder named `traj_segs/` and named following the pattern `iter_XXXXXX.h5`. The basis states are returned to WESTPA in `get_pcoord.sh` as both the “trajectories” of the zero-th iteration and the restart files for propagating the first iteration and recycled walkers. The dynamics is propagated using OpenMM for 100 ps for each iteration, and the output trajectory files and state xml files are returned to WESTPA in `runseg.sh`. These files are deleted once they are returned in order to save disk space. See the sample project setup files for detail.

#### 3.2.4 Running the WE Simulation

Similar to other examples, the simulation can be run using `./run.sh` from the top-level permeability tutorial directory.

#### 3.2.5 Analyzing the WE Simulation

The analysis of the membrane permeation simulation can be found in the accompanying Jupyter notebook titled Membrane Permeability Tutorial (Analysis). In this tutorial notebook, we demonstrate how to extract a complete, continuous pathway of a membrane-crossing event and calculate the incoming flux to the target state from the WE membrane permeation simulation. Note that this tutorial assumes that you already have a completed simulation using the HDF5 framework with at least one crossing event (~40 iterations).

#### 3.2.6 Conclusion

In this tutorial, we have illustrated the relative ease in which one may use the WESTPA 2.0 software package to perform advanced WE path sampling simulations of membrane permeation for a small molecule (butanol). Using a single workstation with two GPUs, our WE simulation can generate membrane permeation events within a few days of wallclock time. WE simulations, when using the WESTPA 2.0 HDF5 framework and MAB binning scheme, are relatively cost effective, both in terms of total compute time and disk storage.

### 3.3 Analysis and restarting with haMSMs: NTL9 Protein Folding

#### 3.3.1 Introduction

Although the WE strategy provides an efficient framework for unbiased rare-event sampling, slow relaxation to steady state and impractically large variance in rate constant estimates may still be limiting factors for complex systems. History-augmented Markov state models (haMSMs) have been demonstrated to provide estimates of steady state from transient, relaxation-phase WE data, which can be used to start new WE simulations [17]. As shown in **Figure 6**, the haMSM plugin for the WESTPA 2.0 software package automatically constructs an haMSM from one or more independent WE simulations to estimate steady-state observables and then can automatically initiate new simulations from those estimates and iteratively repeat this procedure when those simulations complete. The underlying haMSM analysis library, `msm_we`, can also be used to perform stand-alone haMSM analysis of existing WESTPA data.

**Figure 6.**
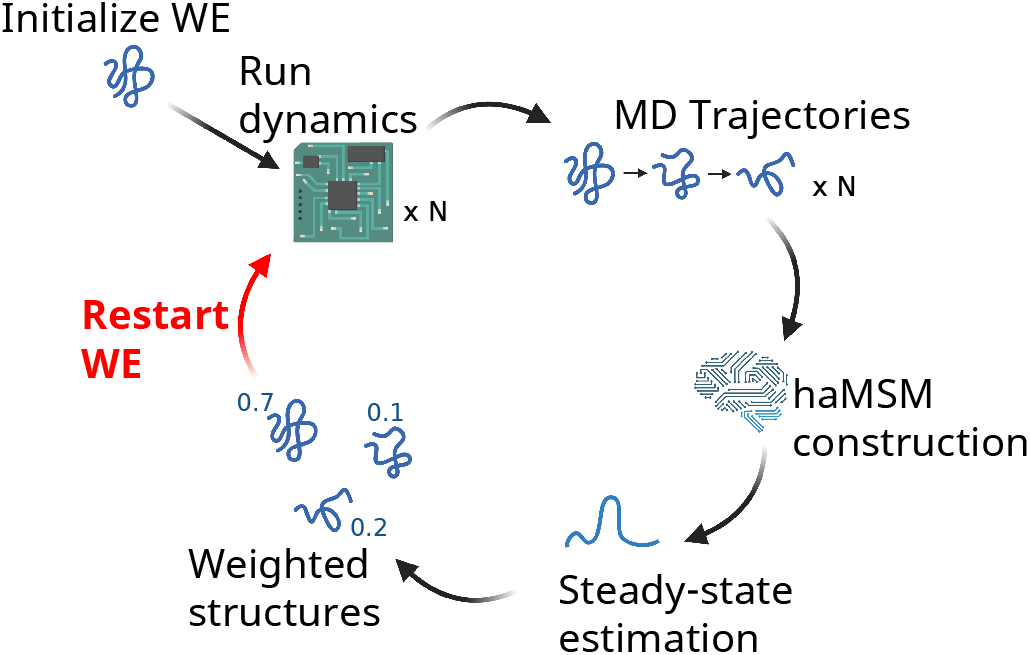
Schematic of haMSM restarting procedure. Trajectories from one or more WE runs are used together to build an haMSM. An estimate of steady-state is obtained from the haMSM, and is used to assign weights to all sampled structures. New WE runs are initiated from these steady-state weighted structures, and the procedure repeats. Figure reprinted with permission from [7]. Further permission related to the source material should be requested from ACS.

##### Learning Objectives

This tutorial demonstrates the use of an haMSM restarting workflow in WE simulations of the ms-timescale folding process of the NTL9 protein. Specific objectives are:

1. How to apply the haMSM plugin for periodic restarting of simulations;
2. How to use the `msm_we` package to build an haMSM from WE data;
3. How to estimate the distribution of first passage times from the haMSM, using `msm_we`.

#### 3.3.2 Prerequisites

The Basic and Intermediate WESTPA Tutorials [32] should be completed before running this tutorial.

##### Computational Requirements

This tutorial can be completed on a computer with a single NVIDIA GTX 1080 GPU and a 2.4GHz Intel Xeon E5-2620 in 90 min. The WE simulation will generate ~5 GB of data, though on a typical cluster filesystem, overhead associated with data redundancy may increase this to ~15 GB. A version of the Amber software package compatible with Amber 16 restart and topology files must be installed to propagate the dynamics and calculate the WE progress coordinate.

The `msm_we` Python package must also be installed, which can be done by first cloning the repository and then installing it into your existing conda environment (with WESTPA already installed) by running:

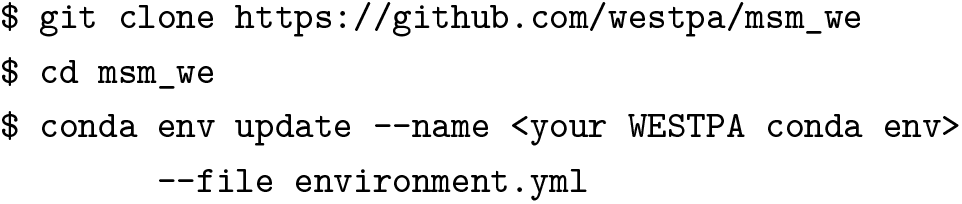

#### 3.3.3 Plugin functionality

Once we initiate a WESTPA run with the haMSM plugin enabled, the plugin will execute a series of independent WE simulations (runs) from the same starting configuration for the number of WE iterations specified in `west.cfg`. For this tutorial, the runs will not use the HDF5 trajectory-saving framework.

If none of the WE runs have reached the target state, the haMSM plugin will sequentially extend each run for a number of iterations specified in `west.cfg`. This extension procedure will be repeated until at least one run has reached the target state. For consistency, all of the other runs in the set will be extended to match the length of this run. As a result of this extension procedure, runs used for the first restart may be longer than runs in subsequent restarts.

After completing the extension procedure, the plugin will construct an haMSM from these runs, and estimate the steady-state distribution and flux into the target state. All structures sampled by the set of runs are used to build this haMSM, and are then weighted according to a steady state. Note that this haMSM uses only the first and last frame of each WE iteration, which effectively sets the lag-time equal to the WE resampling time. A number of plots are automatically generated from the model. The flux profile, shown in **Figure 7**, provides an important metric of convergence, and should be examined carefully. A flatter flux profile indicates more converged weighted ensemble.

**Figure 7.**
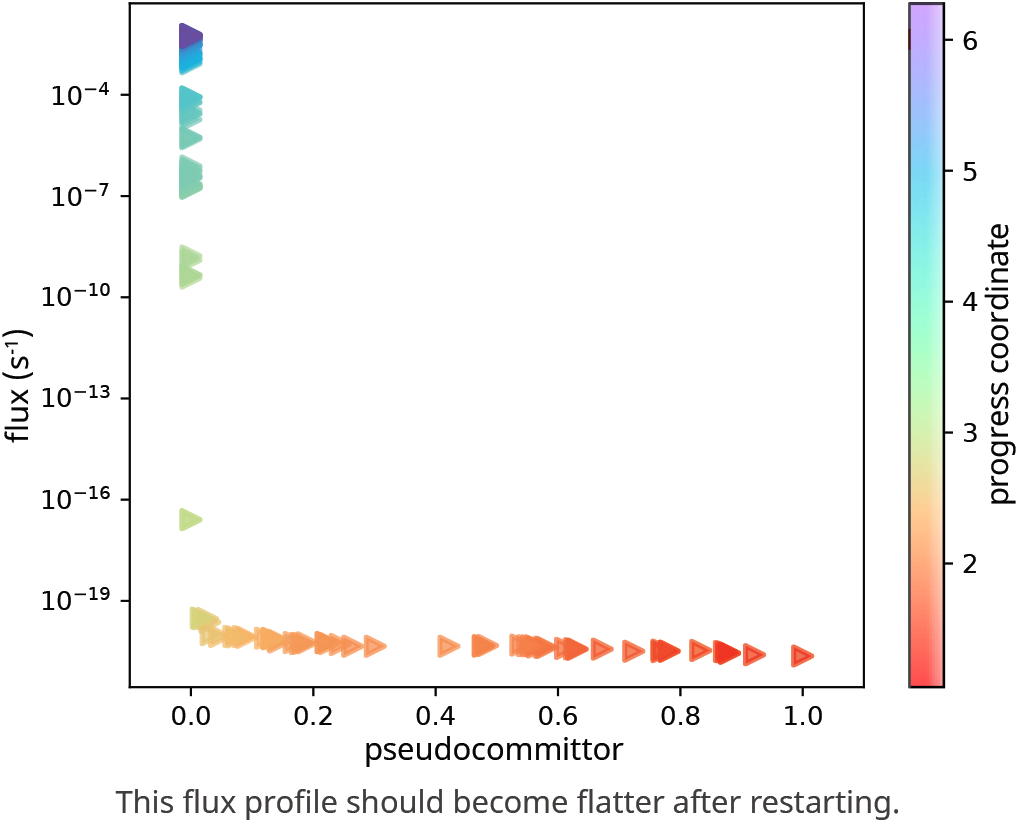
Sample flux profile generated by haMSM restarting plugin after a restart. The x-axis is labeled “pseudocommittor”, since these are committor values generated from a unidirectional path ensemble. This plot should flatten in successive restarts until steady state is reached. The color scale indicates the progress coordinate associated with each pseudocommittor—notably, most of the dynamic range in committor-space is restricted to a small range of progress coordinate values. This image can be found in restart0/plots/psuedocomm-flux_plot_pcoordcolor.pdf after restart is performed.

A new set of runs from the resulting weighted structured are initialized. These runs are correlated but independent from this point onward. As a technical note, when initializing the new WE simulations, these structures are used as “start states”. Within the WESTPA 2.0 framework, start states are a third category of state, in addition to basis states and target states. Like basis states, start states are used for seeding trajectory walkers when initializing a simulation with `w_init`; however, *unlike* basis states, start states are not used after this point and walkers reaching the target state will **not** be recycled into start states but rather only to the basis states. The new WE runs are executed for the number of WE iterations specified in `west.cfg`, in series. At this point, the model is saved, a new restart is prepared, and the process repeats from that point onward.

Please see [7, 17] for more theoretical background on the models used by this plugin.

#### 3.3.4 Preparing the system

##### The system

For our simulation of the NTL9 protein folding process, we use a stochastic Langevin thermostat with low-friction (collision frequency *γ* = 5 ps^−1^) and a generalized Born implicit solvent model. The system consists of ~600 atoms. We will use the haMSM restart plugin to automatically perform three independent WESTPA runs serially before constructing an haMSM. A single restart will be performed from the haMSM steady state estimate, and then WE simulation will be continued for another 106 WE iterations for each of the three runs.

To reduce runtime for this tutorial, we provide a partially completed set of three independent WE simulations. In this set of simulations, the first two runs have completed, and the third is nearing completion. None of these simulations have yet reached the target state.

After continuing this set of WE simulations, the third simulation will finish and reach its maximum number of WE iterations. With very high probability, none will have reached the target folded state, and pre-restart extensions will be triggered. The initialized runs were chosen such that it is unlikely that the third run will reach the target state before the next restart. However, it is possible that in the remaining few WE iterations of the third simulation, this simulation will reach the target state, in which case we will skip the extension procedure.

After a single round of extensions, the target state should be reached in run 2, though not necessarily runs 1 or 3. We will construct the haMSM from the three extended runs, and restart a new set of three runs from the steady-state estimates.

##### Structure of Plugin-Specific Files

The following is a list of some important files used and generated in `$WEST_SIM_ROOT/` by the haMSM plugin.

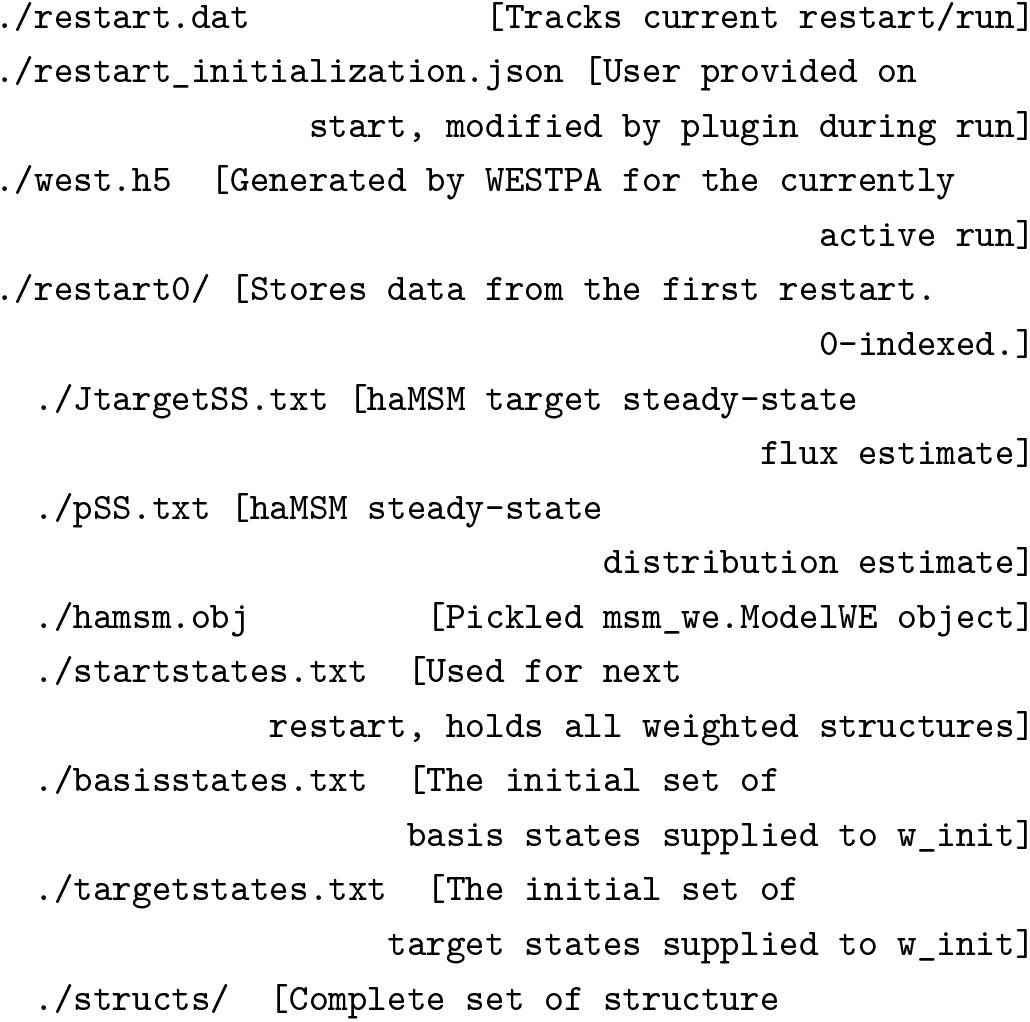
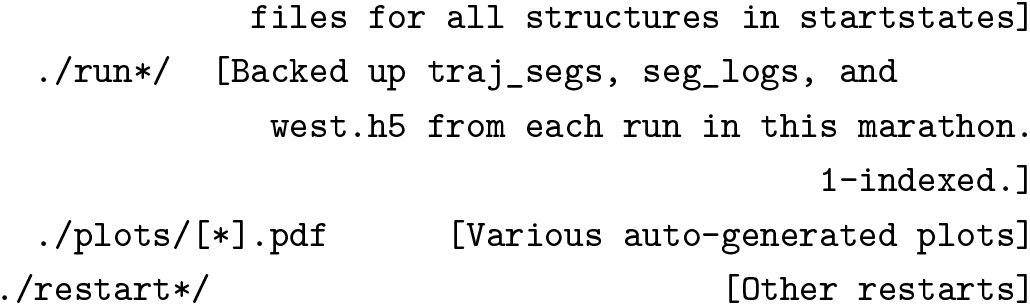

The following files containg more adjustable parameters or are more tightly integrated in the workflow and therefore warrant a more in-depth explanation.

`west.cfg`: The haMSM restarting plugin requires a number of parameters to be set in the appropriate section of `west.cfg`. Details regarding these parameters are listed at https://westpa.readthedocs.io/en/latest/documentation/ext/westpa.westext.hamsm_restarting.html#west-cfg.

`restart_initialization.json`: When initializing each run, the plugin needs to know what configuration it should be launched with. After the first restart, this is automatically generated. However, *before* the first restart (i.e. in producing the initial set of runs in Workflow Step 1), there is no way for the plugin to determine how the first run was initialized. So, the parameters initially passed to `w_init` must be manually entered into `restart_initialization.json`.

`westpa_scripts/restart_overrides.py`: When building the haMSM, some dimensionality reduction is typically necessary as it’s generally neither practical nor useful to analyze the model on the full set of coordinates. This dimensionality reduction is highly system-specific, so no general procedure is distributed with the plugin. Instead, the user is required to define a function which takes in an array of full-atomic coordinates of shape `(n_segments, n_atoms, 3)`, perform the desired dimensionality reduction, and then return the reduced coordinates in an array of shape `(n_segments, n_features)`. This functions are then loaded by the haMSM analysis code at run-time, and used throughout. More details are available at https://westpa.readthedocs.io/en/latest/documentation/ext/westpa.westext.hamsm_restarting.html#featurization-overrides.

##### Preparing the WE Simulation Environment

To prepare the system for using the haMSM restarting plugin, first clone the tutorial repository. In `env.sh`, change `TEMPDIR_ROOT` to point to a directory on your filesystem where temporary files will be created (on a cluster, this should ideally be some node-local scratch/temp space that supports I/O, but on a local workstation can be a new folder such as `$WEST_SIM_ROOT/temp`). This temp space will be used for temporary files created during progress coordinate calculation. In the same file, change `AMBER_EXEC` and `CPPTRAJ` to point to your AMBER and CPPTRAJ executables. In `west.cfg`, change `ray_tempdir` to point to the same directory as `TEMPDIR_ROOT`.

Then examine haMSM plugin-specific configuration files above to familiarize yourself with them, though for this tutorial, no further changes are required. To download and extract the prepared files for the in-progress simulation this tutorial uses, run the following command from within the main simulation directory.

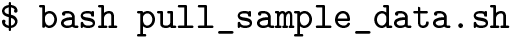

#### 3.3.5 Running the WE simulation

Now, we are ready to (re)start the WE simulation. The haMSM plugin will automatically perform the restarting and analysis. Typically, we would initialize the system using `w_init` as we do for all WE simulations using the WESTPA 2.0 software package. However, to reduce the runtime for this tutorial, we have provided a pre-prepared system, and running `w_init` is not required. Once we have configured the haMSM plugin through `west.cfg`, we can restart the WE simulation and run the simulation for a few iterations, by simply executing the following command.

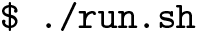

If the target state has not been reached after running for the specified number of iterations, additional rounds of restarting/extending the WE simulation will automatically be launched. Once the target state has been reached, the plugin will build an haMSM, update statistical weights for each sampled structure, and restart a new set of WE trajectories initialized from those structures with updated weights. This will all be done automatically.

##### Start States

After performing a restart, we will find under the `restart0/` directory a `startstates.txt` reference file that lists all the structures used for the restart (start states) and their associated weights for initializing new WE simulations from the first round of restarting. As noted, these are distinct from the basis states. The `startstates.txt` text file is formatted with three columns, which define the names (e.g., “b21s0”), associated probabilities, and name of the directory containing the structure files of the start states (relative to the path defined under `west.data_refs.basis_states` in `west.cfg`). Structure files corresponding to these start states are in `restart0/structs/` and are named according to the structure’s haMSM bin and the structure’s index within that bin. Start states can be added to the pool of potential structures for WE initialization by adding the `--sstates-from` or `--sstates` option to the `w_init` command. Similar to the `--bstates` options for basis states in the above **Advanced Tutorial 3.1**, the `--sstates-from` option is used to indicate a text file with a list of start states and the `--sstates` option is used to append additional start states through the command line.

#### 3.3.6 Analyzing the WE simulation

After the plugin finishes running, you will find the associated `west.h5` for each run and the associated haMSM pickled `hamsm.obj` objects for each marathon in the `restart*/` directories. Although the plugin will automatically build the haMSMs and perform some of the analysis based on the configuration files, haMSM analysis can also be manually performed post-simulation on WESTPA data with the `msm_we` library (as used internally by the plugin).

For this tutorial, you can use either the data generated by the steps above, or the pre-prepared `west.h5` files containing data generated from a similar simulation configuration. This analysis largely follows the `msm_we` usage instructions provided in the `msm_we` documentation (https://jdrusso.github.io/msm_we/usage.html).

For detailed instructions on how to analyze your simulation results, please refer to the Jupyter notebook distributed along with this tutorial.

#### 3.3.7 Conclusion

Complex systems may exhibit relaxation slow enough to prevent direct measurement of rate constants using probability flux in WE. This tutorial therefore presents the haMSM plugin for leveraging relaxation-phase WE simulations by automatically (i) building a haMSM; (ii) generating an estimate of the steady-state probability distribution and the corresponding steady flux; and (iii) if desired, restarting a new WE simulation or set of simulations from the estimated steady state. Each haMSM yields an estimate for the MFPT and FPT distribution using `msm_we`.

### 3.4 Creating Custom Analysis Routines and Calculating Rate Constants

#### 3.4.1 Introduction

In this tutorial, we will go over how to create custom analysis routines using the `westpa.analysis` Python API and how to calculate rate constants using the Rate from Event Durations (RED) analysis scheme, which enables rate-constant estimates from transient, presteady-state data and therefore shares the same motivation as the haMSM analysis scheme [7, 17] used in the above **Advanced Tutorial 3.3**. For the creation of custom analysis routines, we will focus on the membrane permeability simulations completed in **Advanced Tutorial 3.2**. For the calculation of rate constants, we will focus on previously published protein-protein binding simulations involving the barnase/barstar system [4].

##### Learning objectives

Specific learning objectives for this tutorial include:

1. How to access simulation data in a `west.h5` file using the high-level `Run` interface of the `westpa.analysis` Python API and how to retrieve trajectory data using the `BasicMDTrajectory` and `HDF5MDTrajectory` readers;
2. How to access steady-state populations and fluxes from the `assign.h5` and `direct.h5` data files, convert fluxes to rate constants, and plot the rate constants using an appropriate averaging scheme;
3. How to apply the RED analysis scheme to estimate rate constants from shorter trajectories.

#### 3.4.2 Prerequisites

In addition to completing the Basic and Intermediate WESTPA tutorials [32], a prerequisite to this tutorial is completion of the above **Advanced Tutorials 3.1 and 3.2**.

##### Computational requirements

Users should have access to at least 1 CPU core for running the analysis tools. For larger datasets, one may want to parallelize some of the tools (especially `w_direct`). The size of a dataset is mainly determined by the number of iterations. For a dataset of greater than 1000 iterations, it may be best to use at least 4 CPU cores at a time.

#### 3.4.3 Creating custom analysis routines

For this part of the tutorial, we will create custom analysis routines using the `westpa.analysis` API for the membrane permeability simulations completed in **Advanced Tutorial 3.2**.

The main abstraction of the `westpa.analysis` API is the `Run` class, which provides a read-only view of the data in the main WESTPA output file (`west.h5`). We will start by opening the `west.h5` file from the permeability run. We assume that the current working directory is the simulation root directory you are interested in analyzing, though the resulting `west.h5` file from **Advanced Tutorial 3.2** is linked to the main directory of this tutorial for convenience. Open a Python interpreter and run the following commands.

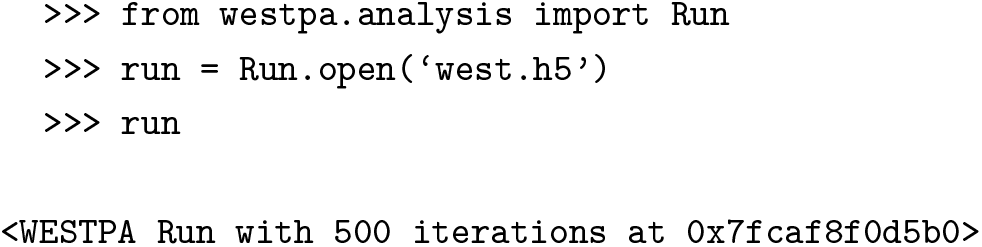

We now have convenient access to a wealth of information about the permeability simulation, including all trajectory segments at each WE iteration and any data associated with those segments, including values of the progress coordinate and other auxiliary data. Iterating over a run yields a sequence of `Iteration` objects, each of which is a collection of `Walker` objects. For example, the following loop iterates over all trajectory walkers in a run, but does nothing with each trajectory walker:

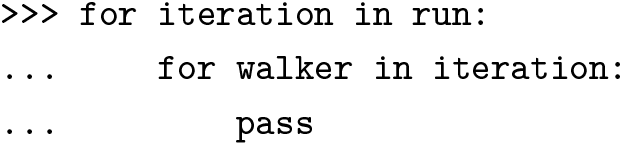

We can access a walker by providing its (1-based) iteration number and (0-based) segment ID:

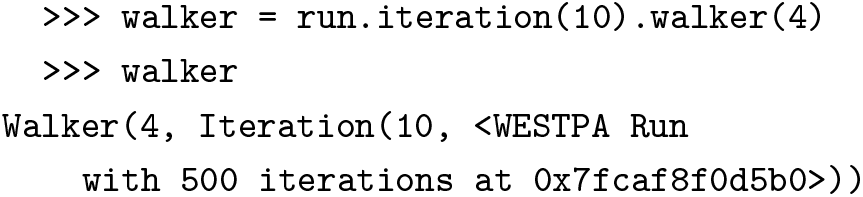

To access the progress coordinates of a certain trajectory walker, we use the pcoords attribute:

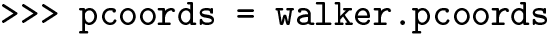

Other properties available through this Python API include the weight, parent and children of a trajectory walker. We can access auxiliary data by looking up the dataset of interest in the `auxiliary_data` dictionary attribute (note that the following auxiliary dataset is not actually present, and the command is provided as an example):

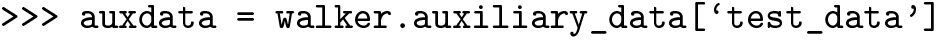

We can also view a list of all recycled (successful) trajectory walkers and choose one walker to trace its pathway through the membrane:

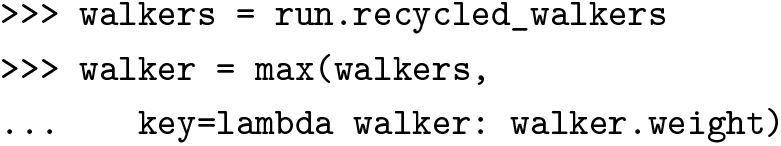

The history of a trajectory walker can be traced by using the `trace()` method, which returns a `Trace` object:

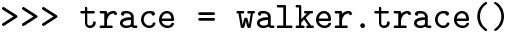

Using the WE iteration and IDs of the trajectory segments obtained from this trace, we can plot the data of our traced trajectory to see how that property is changing. Remember that the `test_data` auxiliary dataset does not actually exist, but can be replaced with an auxiliary dataset of your choice.

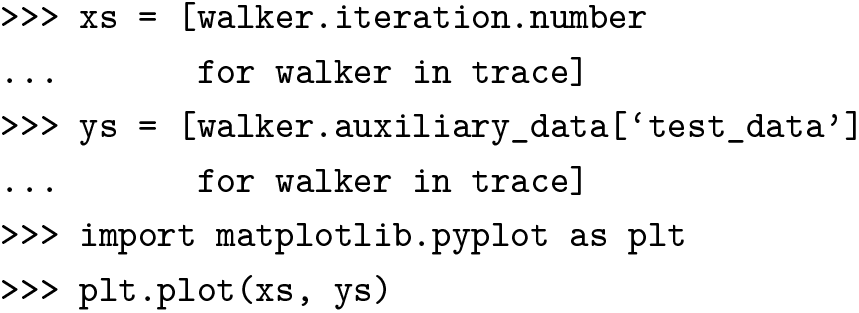

One goal of the `westpa.analysis` API is to simplify the retrieval of trajectory coordinates. Two built-in readers are provided for retrieving MD trajectory coordinates: (1) `BasicMDTrajectory`, which handles trajectory files outputted by the dynamics engine; or (2) `HDF5MDTrajectory`, which handles trajectories stored using the new HDF5 framework, as is done in the above **Tutorial 3.1 and 3.2**. For users requiring greater flexibility, custom trajectory readers can be implemented using the more general `Trajectory` class. Here we provide a brief overview of both the `BasicMDTrajectory` and the `HDF5MDTrajectory` readers. The following is included only as an example, since the trajectory files required are not provided. MD trajectories stored in the traditional manner may be retrieved using the `BasicMDTrajectory` reader with its default settings:

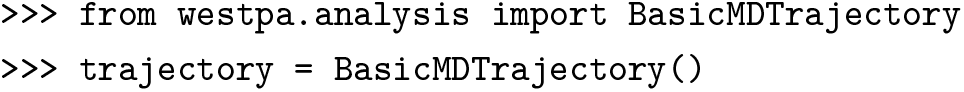

Here, `trajectory` is a callable object that takes either a `walker()` or a `trace()` object as input and returns an `mdtraj.Trajectory()` object (https://mdtraj.org/1.9.5/api/generated/mdtraj.Trajectory.html). To retrieve the trajectory of the trace obtained above, then save the coordinates to a DCD file (e.g., for visualization using the VMD program), we can run the following:

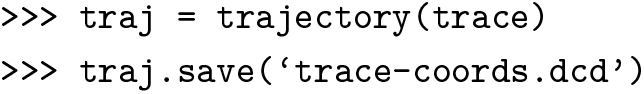

Note that in the code above, we have relied on the fact that the `traj_segs/` directory of interest is contained in the current working directory. In the general case, the name of the simulation root directory may be provided to the trajectory reader via the optional `sim_root` parameter. Minor variations of the “basic” trajectory storage protocol (e.g., use of different file formats) can be handled by changing the parameters of the `BasicMDTrajectory` reader:

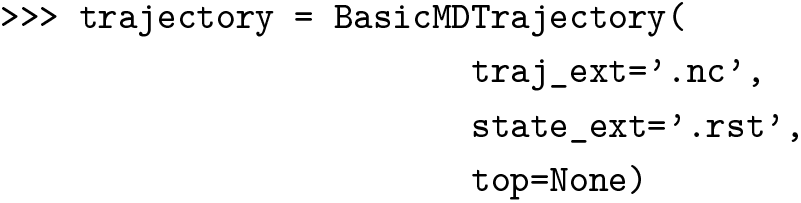

However, suppose that instead of storing the coordinate and topology data for trajectory segments in separate files (`seg.dcd` and `bstate.pdb`), we store them together in an HDF5 trajectory file (such as `iter_XXXXXX.h5`) using the new HDF5 restarting framework available in WESTPA 2.0. This change can be accommodated by using the `HDF5MDTrajectory` reader:

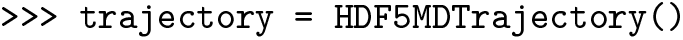

The examples above highlight the flexibility and convenience provided by the `westpa.analysis` package and provide the building blocks available to a user wanting to explore the `west.h5` file and create custom analysis routines using data in the `west.h5` file.

#### 3.4.4 Calculating rate constants using the original WE scheme

For the two remaining sections of this tutorial, we will focus on applying the RED analysis scheme [19] to calculate the association rate constant from previously published proteinprotein binding simulations involving the barnase/barstar system [4]. The RED scheme involves three steps:

##### 1. Calculate State Populations and the Flux into the Target State

The target state can be defined using either the WE progress coordinate or auxiliary coordinates. The analysis bins and state definitions are placed in the analysis section of the `west.cfg` file.

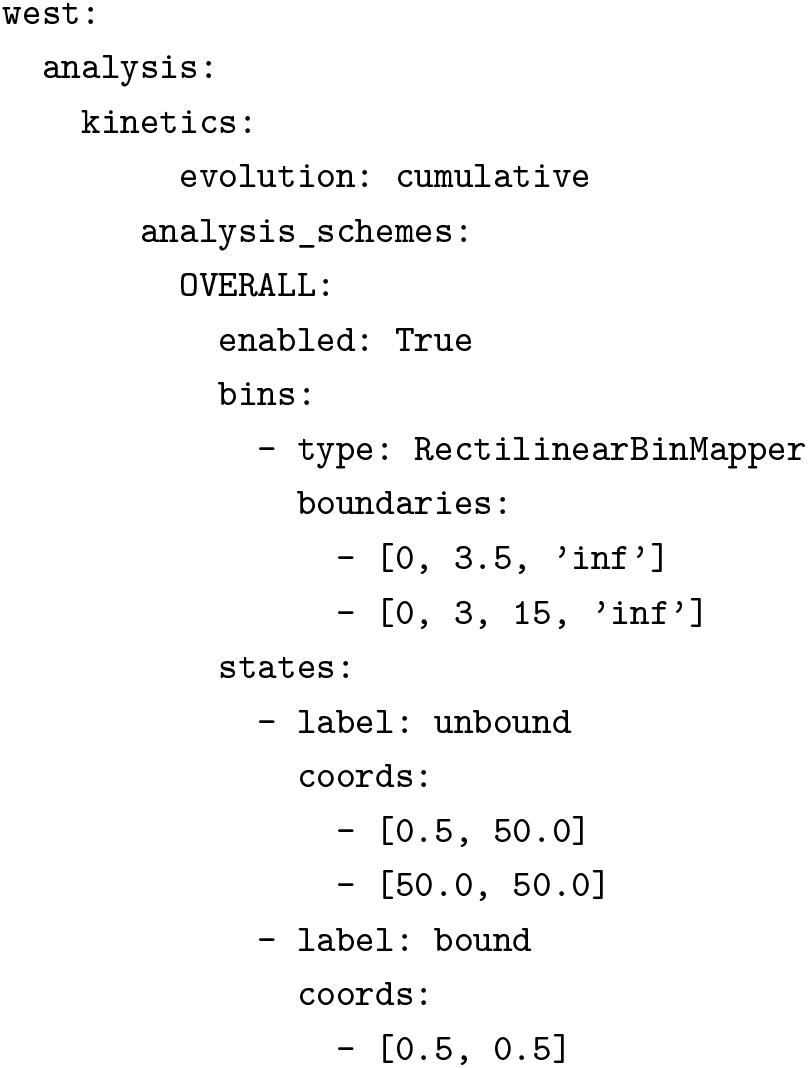

To calculate the flux into a state defined by the progress coordinate, we can use the `w_ipa` program. Flux calculation with `w_ipa` involves two steps: (1) assigning trajectory segments to states using the `w_assign` command-line tool, and (2) calculating the probability flux between each pair of defined states using the `w_direct` command-line tool. Given the large size of the barnase-barstar simulation HDF5 file, we have provided the resulting `assign.h5` and `direct.h5` files for the remainder of the tutorial. To use `w_ipa` for flux analysis, one would run the following at the command line:

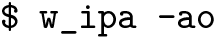

This command will analyze the `pcoord` data from `west.h5` using the scheme for bins and states defined in `west.cfg` and not drop the user into an ipython environment (to make use of that functionality, remove the `-ao` options from the above command). The resulting `assign.h5` and `direct.h5` files, the latter containing the fluxes, will be outputted to a newly created directory that is named for the relevant scheme (in this case that will be in `./ANALYSIS/OVERALL/`. The `evolution:cumulative` option (which is the default option) ensures that all evolution datasets are calculated with a rolling average, a requirement for using the RED scheme (see below).

To calculate the flux into a state defined by auxiliary coordinates, we still use the scheme for bins and states defined in the `west.cfg` file. However, instead of using the `w_ipa` program, we use the command-line tools, `w_assign` and w_direct. Before using these tools, we need to copy `module.py` to your current directory first (by default, `module.py` is located in the `./scripts/` directory). Then, we can assign trajectory segments to specified states using the `w_assign` tool:

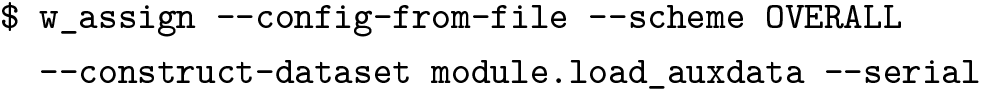

The `--config-from-file` option tells the program to read analysis parameters from the `west.cfg` file’s analysis section and the `--scheme` option specifies the relevant scheme for bins and states. The `--construct-dataset` option provides a function to `w_assign` for loading in the auxiliary data which is located in the file `module.py`. The `--serial` option tells `w_assign` to run the assignment in serial mode. Running `w_assign` will generate an `ANALYSIS/` directory and place an `assign.h5` file in a scheme-specific folder there. Next, we apply the `w_direct` tool to the `assign.h5` file.

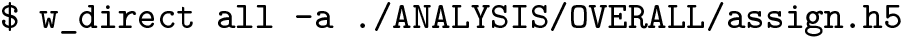

This command will generate a `direct.h5` file in the same directory where we ran the `w_direct` command. Move this file to the `analysis/` folder that was generated by `w_assign` and proceed with the analysis. Note that the above is not part of the following tutorial and is only included to provide an example to users of how to perform an analysis on auxiliary data.

##### 2. Correct the Fluxes using the RED Scheme

To correct the calculated fluxes using the RED scheme, we apply the `w_red` command-line tool, which will read in the `rate_evolution` and durations datasets in your `direct.h5` file and calculates a correction factor for the flux value at each iteration. To use this tool, add the following to the analysis section of your `west.cfg` file:

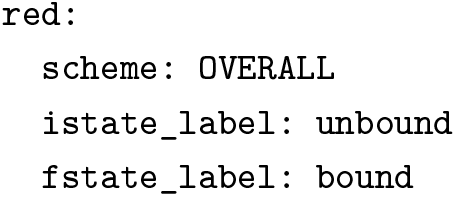
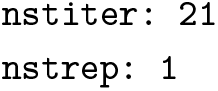

For the scheme option, specify the desired scheme for bins and states to use from your `west.cfg` analysis section and for the `istate_label` and `fstate_label` options, specify the initial and target states, respectively. The `nstiter` parameter is the number of frames per WE iteration that were saved during the WESTPA simulation and `nstrep` is the frequency of outputting progress of the RED calculation. After setting all parameters, run the `w_red` tool from the command line:

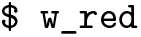

A new dataset containing the corrected fluxes, named `red_flux_evolution`, will be added to your `direct.h5` file. If that dataset already exists, the `w_red` tool will ask if you want to overwrite the existing dataset.

The new `red_flux_evolution` dataset is created by adjusting the `rate_evolution` dataset by the cor-rection factor determined by the RED scheme. The `rate_evolution` dataset, in turn, is composed of the relevant `conditional_flux_evolution` dataset (from `direct.h5`) normalized by the steady state populations from `labeled_populations` (from `assign.h5`). Therefore, when using the RED scheme (or the original `rate_evolution` dataset), no explicit normalization of the fluxes by the steady state populations is necessary.

##### Convert the Flux to a Rate Constant

The fluxes that we just calculated are already rate constants in units of inverse *τ* (the resampling interval used for your WE simulation).

For a unimolecular process, the final units of the rate constant should be inverse time (e.g., s^−1^) and can be obtained by converting from units of per *τ* for the extracted flux array to the desired time unit. Note that the *τ* value needs to be provided by the user and is not currently recorded in the `west.h5` file.

For a bimolecular process, which is the case for our `OVERALL` scheme, the rate constant should be in units of inverse time and inverse concentration (e.g., M^−1^s^−1^) and can be obtained by first converting to the desired time unit, as done for unimolecular processes, and dividing the resulting flux values by the effective concentration of the solutes involved in the bimolecular process given the volume of the simulation box. In the case of our barnase-barstar system, the *τ* value was 20 ps and the effective concentration for the barstar ligand was 1.7 mM. We will therefore divide all of the `conditional_flux` values by 20 × 10^−12^ s and 0.0017 M to obtain per iteration rate constants in units of M^−1^s^−1^.

#### 3.4.5 Monitoring Convergence of the Rate Constant

To monitor the convergence of the rate-constant estimate, we can plot the time-evolution of the rate constant using both the original and RED schemes and assess how close our original estimate is to the RED estimate. If the two schemes are converging to the same value, that can be one indication that the simulation has begun to converge.

To obtain the rate constant using the original scheme [1], simply convert the `rate_evolution` dataset to the appropriate units as discussed above. The time-evolution of this rate-constant estimate can then be compared with the RED-scheme estimate to assess convergence of the simulation to a steady state (**Figure 8A**).

**Figure 8.**
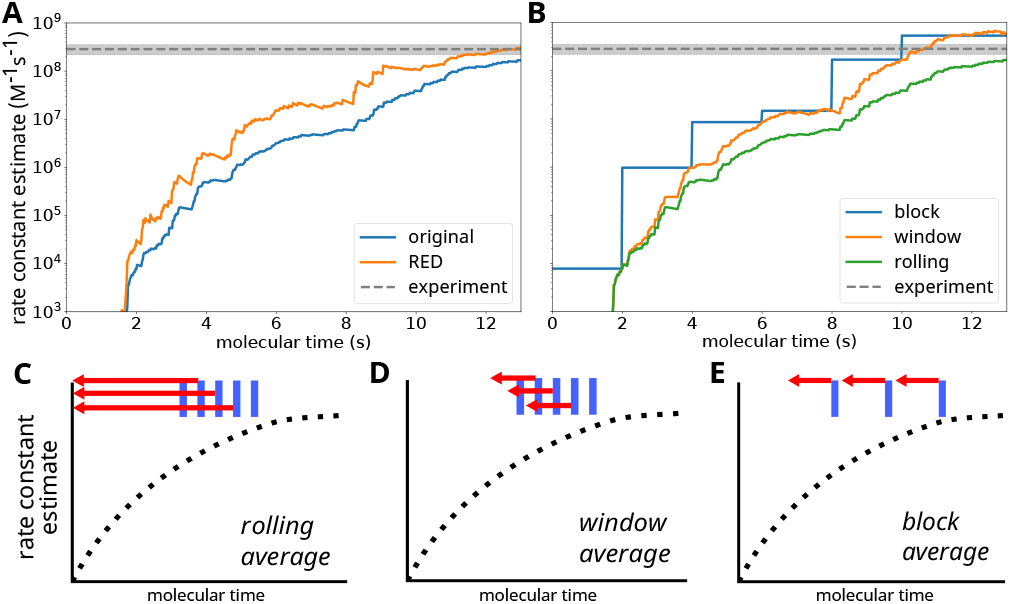
A) A comparison of the RED and original schemes for estimating the rate constant for protein-protein association involving the barnase/barstar system from previously published WE simulations [4]. In this case, the simulation has not converged to a steady state, as indicated by different rate-constant estimates using the original and RED schemes. B) A comparison of different averaging schemes using the same dataset. While the RED scheme is, in principle, only compatible with rolling averages, the use of block and window averaging can still be informative for monitoring convergence of the simulation. C-E) Schematic illustrations of rolling, window, and block averaging schemes, indicating points where averages are calculated (blue) and the extent of data used for the averages (red).

##### A note on averaging schemes

There are three main types of averaging schemes that can be used to monitor convergence when plotting the rate constant evolution using the original scheme. It may be useful to plot different schemes depending on the behavior of your specific system. A few examples are shown here with instructions on how to generate the plots.

The first averaging scheme is the default which is a rolling average (**Figure 8B**), which can be achieved by specifying `evolution: cumulative` in the analysis section of `west.cfg` and setting `step_iter: 1`. This method of visualizing the rate constant evolution offers the smoothest curve and is recommended as the most stringent way of assessing convergence, as it incorporates information from the entire simulation. A rolling average is also implicitly incorporated into the RED scheme, which by design never excludes data from the start of a simulation. When analyzing rate constant estimates generated by the RED scheme, specify `evolution: cumulative` to ensure that only the implicit rolling average is performed.

The second averaging scheme is a window average, which can be achieved in `w_ipa` by specifying `evolution: cumulative` and `step_iter: 10`, or whatever your desired averaging window is. A recommended starting averaging window size is 10% of the length of your simulation, but the most robust would be the lag time of your simulation as determined from an autocorrelation of the flux plot. A windowed average is not as smooth as the rolling average but can give a better indication of convergence at different stages of your simulation relative to other stages.

The third and final averaging scheme is a block average. This will require setting `evolution: blocked` and `step_iter: 10`. The rationale behind choosing the block size here is the same as the window size discussed above. The block average will appear like a step function where each block an average of the preceding block is plotted. This method of plotting is the least smooth, but can be best for obtaining a final value of the rate constant that is not influenced by earlier, lower values.

#### 3.4.6 Conclusion

Among the upgrades introduced in the WESTPA 2.0 software package are ones that enable the creation and execution of more streamlined analysis of simulations and more efficient estimation of rate constants. The `westpa.analysis` subpackage can be utilized to more carefully inspect WESTPA trajectory data and to create custom analysis routines. The RED analysis scheme for correcting rate constants based on the “ramp-up” time in the fluxes is implemented in the `w_red` command-line tool. The files contained in this tutorial for utilizing the RED scheme are intended to provide useful starting points for analyzing the kinetics of WESTPA simulations.

### 3.5 M-WEM Simulations of Alanine Dipeptide

#### 3.5.1 Introduction

The Markovian Weighted Ensemble Milestoning (M-WEM) approach [10] is a modified version of the Weighted Ensemble Milestoning (WEM) [40, 41] approach. Both approaches are designed to use the WE strategy to enhance the efficiency of the milestoning method in calculating equilibrium and nonequilibrium properties (e.g., free energy landscape and rate constants, respectively).

In the milestoning method [11, 12] the reaction coordinate is stratified using multiple high-dimensional interfaces— or milestones. Short trajectories are propagated between the interfaces and using the principles of continuous time Markov chains, the properties of a long timescale process can be calculated. But the milestones need to be placed significantly far from each other to lose memory of the previous milestone. But converging the transition statistics between distant milestones can be expensive depending on the underlying free energy landscape and the complexity of the system. Because of the use of shorter trajectories that do not require trajectories to transit from the initial to final states, the WEM and M-WEM calculations are computationally less expensive than a WE simulation. On the downside, however, one cannot obtain continuous pathways due to the lack of continuous trajectories between the starting and the final state.

In the M-WEM approach, regions between the milestones are referred to as “cells”. WE simulation is performed within the cell with half-harmonic walls present at the milestone interfaces to prevent trajectory escape [10, 42, 43]. In this tutorial, we will use M-WEM to calculate the mean first passage time (MFPT), free energy landscape and committor function for the conformational transition in the alanine dipeptide system.

##### Learning objectives

This tutorial covers the installation of and use of the Markovian Weighted Ensemble Milestoning (M-WEM) software in combination with WESTPA to compute the kinetics and the free energy landscape of an alanine dipeptide. Specific learning objectives include:

1. How to install the M-WEM software and perform an M-WEM simulation;
2. How to create milestones to define the M-WEM progress coordinate;
3. How to analyze an M-WEM simulation to compute the mean first passage time, committor, and free energy landscape.

#### 3.5.2 Prerequisites

The users should have a basic understanding of running WE simulations using the Minimal Adaptive Binning (MAB) scheme, and should have completed **Advanced Tutorial 3.2** before commencing the M-WEM tutorial. Also, a basic idea of the Markovian Milestoning framework is necessary. For that purpose, the users should refer to the following manuscripts [10, 42, 43].

##### Computational requirements

In terms of software, this tutorial requires several Python modules (Numpy, Scipy, and Matplotlib) in addition to the WESTPA 2.0 software and NAMD 2.14 simulation package. Note: M-WEM is implemented using the colvars module in NAMD. Please check out the NAMD tutorial (http://www.ks.uiuc.edu/Training/Tutorials/namd-index.html) and colvars tutorial (https://colvars.github.io/colvars-refman-namd/colvars-refman-namd.html).

In terms of computer hardware, this tutorial will require approximately 4 GB of disk space. Running the simulation 100 WE iterations for each milestone takes ~85 min on an Intel Core i5-8250U CPU @ 1.60GHz processor with 4 CPU cores. For 8 milestones that amounts to about 11-12 hr if the calculation is performed serially with 4 CPU cores used at a time. But if a computer cluster is available, each milestone should be run in parallel which will significantly reduce the wall clock time. The analysis for each milestone takes ~5 min for each milestone with the same computing hardware but with 1 CPU core. For 8 milestone cells that would be ~40 min, but similar to the simulation, the analysis for each milestone can be done in parallel.

#### 3.5.3 Installation of the M-WEM software package

The `M-WEM` software package can be downloaded from https://github.com/dhimanray/MWEM. To install the package go to the main directory that contains the `setup.py` file and run the following command:

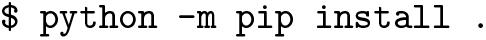

The M-WEM software should be installed in the same conda environment in which WESTPA 2.0 is installed.

#### 3.5.4 Setting up a M-WEM environment

##### Overview

For performing the M-WEM simulation, we need to first create the milestone anchors along the transition pathway. This is a typical prerequisite for milestoning simulation. It is typically done using steered MD simulation [44] which is a common technique in MD simulation. To avoid spending extra time and possible variability in the results, we have generated the milestone anchors and provided them in the anchors directory.

##### The system

We will be studying the conformational change of gas phase alanine dipeptide using the M-WEM scheme. The details of this example are provided in the article: Ray et al. 2022 [10].

The free energy landscape for the system is shown in **Figure 9**. To simulate the transition from state A to state B, 9 milestones are placed at *ϕ* = −80^*◦*^, −60^*◦*^, −40^*◦*^, −20^*◦*^, 0^*◦*^, 20^*◦*^, 40^*◦*^, 60^*◦*^, 80^*◦*^. This created 8 cells bound by the milestones. The anchors are chosen in a way that they are located approximately in the middle of each cell.

**Figure 9.**
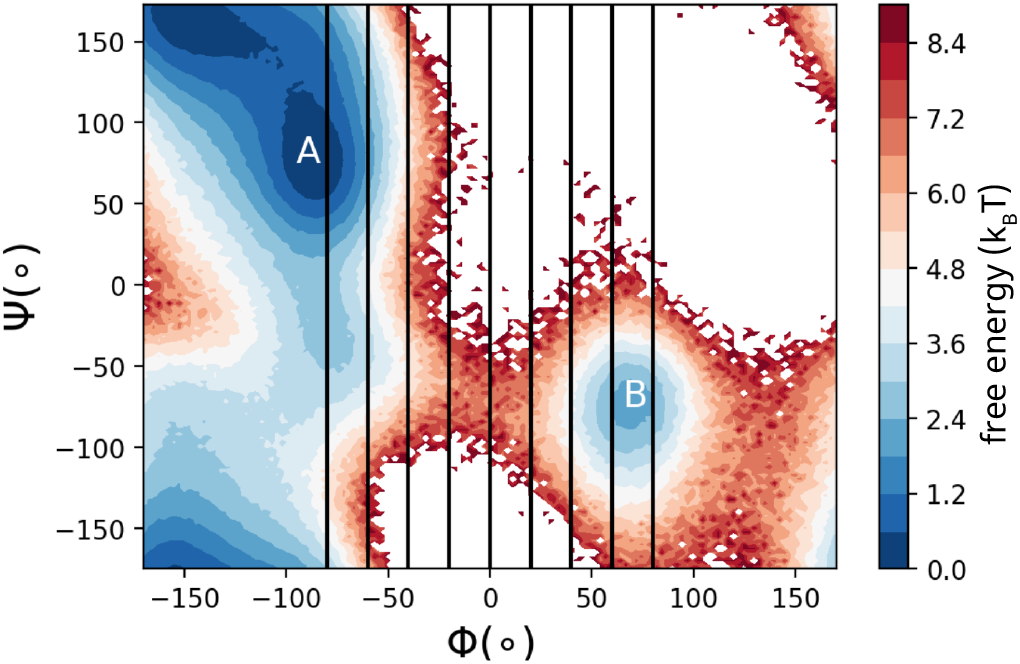
Free energy landscape of alanine dipeptide in the gas phase. Milestone positions are shown as vertical lines. Our aim is to simulate the MFPT of the transition from state A to state B.

##### Preparing the simulation environment

The `tutorial-3.5/` directory contains all simulation and analysis files and will be referred to as the simulation home directory for the rest of the tutorial. The milestone anchors for all cells generated from the steered MD simulation are provided as pdb files in the `anchors/` directory.

The `build.py` script is a python script which will set up all the milestones for the simulation. Each milestone cell will be simulated in a different directory, numbered from 0 to 7.

The `template/` directory is a generic template for M-WEM simulation for any one cell. It contains all necessary files except for the pdb files which are specific to each cell (which are in the `anchors/` directory). Also, the `colvars.in` files have replaceable strings which are used by the `build.py` script to create cell specific files. For example, there are terms like “CENTER”, “HIGH”, and “LOW”. These are places where the position of the center and the two milestones are written by the `build.py` code. Make sure to edit the `env.sh` file to include the path to your NAMD installation.

In the `common_files/` directory the topology (.psf), the parameter (.prm) and the NAMD configuration files are provided. The structure file (.pdb) in this directory and in the `equilibration/` directory are prepared by the `build.py` script, and are different for each milestone cell.

Once you have your M-WEM setup ready, prepare the cell-specific folders by running the following command:

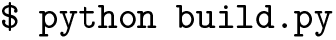

#### 3.5.5 Running the M-WEM simulation

In the `equilibration/` directory of each cell, a constrained equilibration of the anchor will need to be performed to keep the anchor approximately in the middle of the cell. The `milestone_equilibration.colvars.traj` file contains the collective variable information, which, in this case, are the Phi and Psi torsion angles of the alanine dipeptide. The `milestone_equilibration.xsc, milestone_equilibration.coor` and `milestone_equilibration.vel` files are NAMD restart files that will be used to start WE trajectories from the endpoint of the equilibration simulation.

The running of the M-WEM simulation for each individual cell is done separately, for the convenience of parallelizing it in a computing cluster. The `run.sh` script performs the simulation via the command `./run.sh`. Unlike typical WESTPA setups, here the initialization setup code is included in the `run.sh` script. Although it does not make a significant difference for a small system like this, we found it is more convenient to submit multiple milestone jobs to a computer cluster by using a single `run.sh` script. Both the equilibration and running of each cell is performed by executing the following command from within each cell-specific folder:

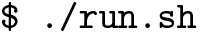

In the first part of the script, equilibration is performed. Then relevant files are copied to the `bstates/` directory, from which they are read by the `westpa_scripts/runseg.sh`. Then the WE simulation is initialized and propagated as usual by the `w_init` and `w_run` commands. In this example script, only one trajectory is propagated at a time. But this can be parallelized based on the computing resources available. Alternative Slurm scripts for running and restarting the simulation are also provided in the same directory.

The total number of iterations performed per milestone is 100. The user may choose to change this number according to their preference. The results reported in this work are from 100 iterations. The convergence is achieved after 40 iterations in our calculation. But it may slightly vary for independent calculations.

#### 3.5.6 Analyzing the M-WEM simulations

After the M-WEM simulations are completed, it is important to properly analyze the results. Please refer to the M-WEM publication for the theoretical details of the analysis [10]. We perform the analysis in two steps:

##### Step 1

Move into the `analysis/` directory and execute the following command:

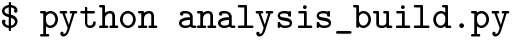

The `analysis_build.py` script will produce directories `cell_0/` through `cell_7/` and copy the corresponding `west.h5` files (WESTPA output files) from the `propagation/` directory into each cell. It will also copy the `west.cfg` files (different from the `west.cfg` files for propagation), and the `analysis.py` files from `westpa_analysis_files/` directory to each cell. The `analysis.py` file also has strings like “LOW” and “HIGH”, which will be replaced by floating point numbers corresponding to the left and right milestones.

Running the `analysis.py` script from within a specific cell directory will produce the `trajectories.pkl`, `crossings.pkl` and `weights.txt` files. The files generated by `analysis.py` contain information on the trajectory traces (history of the segments in the final iteration), the time and location (which milestone right or left) of the milestone crossings, and the weights of each traced trajectory respectively.

Perform analysis in all cells by running the following command from within the `analysis/` directory:

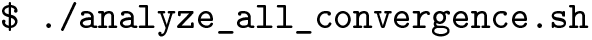

This script will execute `python analysis.py` from within each cell-specific directory to produce the .pkl and .txt outputs for the final iteration. But, for the sake of checking convergence of our results, it will also produce similar files for some subsequent iterations. To do that, the script will copy the `analysis.py` to `analysis_x.py` (where x = iteration number) and replace the `w.niters` inside each `analysis.py` to the corresponding iteration number. Then, it will produce `trajectories_x.pkl`, `crossings_x.pkl` and the `weights_x.txt` files for each x. This step can take several minutes to a few hours depending on the computing hardware. If you have access to a computing cluster, you may choose to submit this as a job. Note that the `analyze_all_convergence.sh` script is customizable. For example, if you want to run all cells in parallel on a cluster you can create separate bash scripts for each cell. Also, `analysis_build.py` will produce the following directories for milestoning analysis in Step 2: `cell_probability/, N_i_j_files/, R_i_files/,` and `committor/`.

##### Step 2

After the analysis of the WESTPA output files are done, we will proceed to analyze our results using the Markovian milestoning framework in two Jupyter notebooks: `kinetics.ipynb` and `free-energy-landscape.ipynb`.

First run the `kinetics.ipynb` notebook to obtain the mean first passage time and the committors. This will also produce the probability distribution file in the milestone space. Details can be found inside the notebook. The MFPT convergence plots and the committor functions should look like **Figures 10 and 11**.

**Figure 10.**
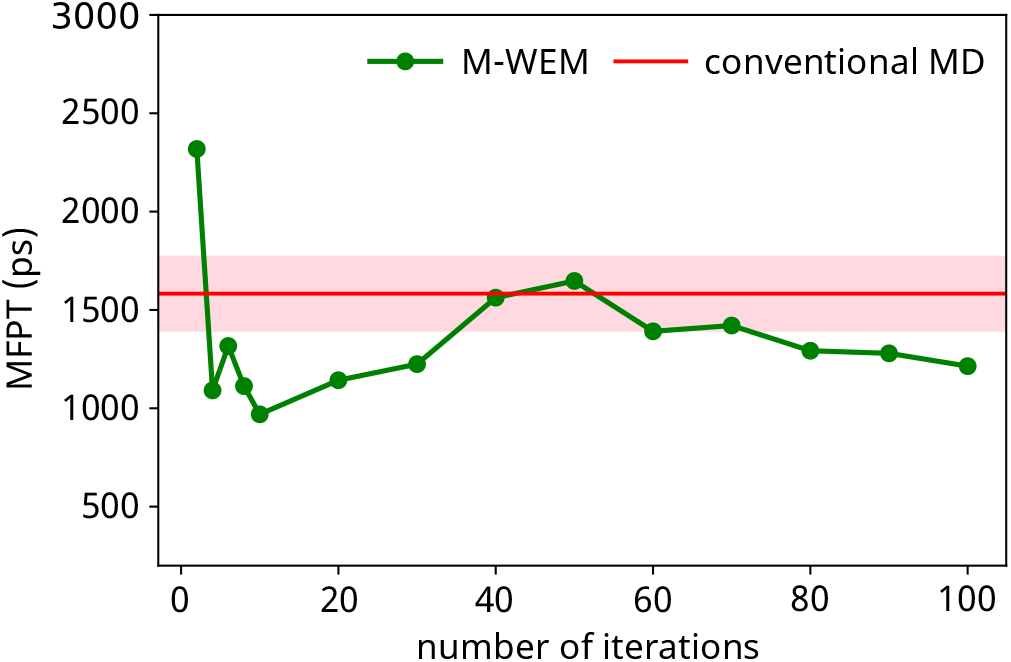
Convergence of the mean first passage time (MFPT) as a function of M-WEM iterations.

**Figure 11.**
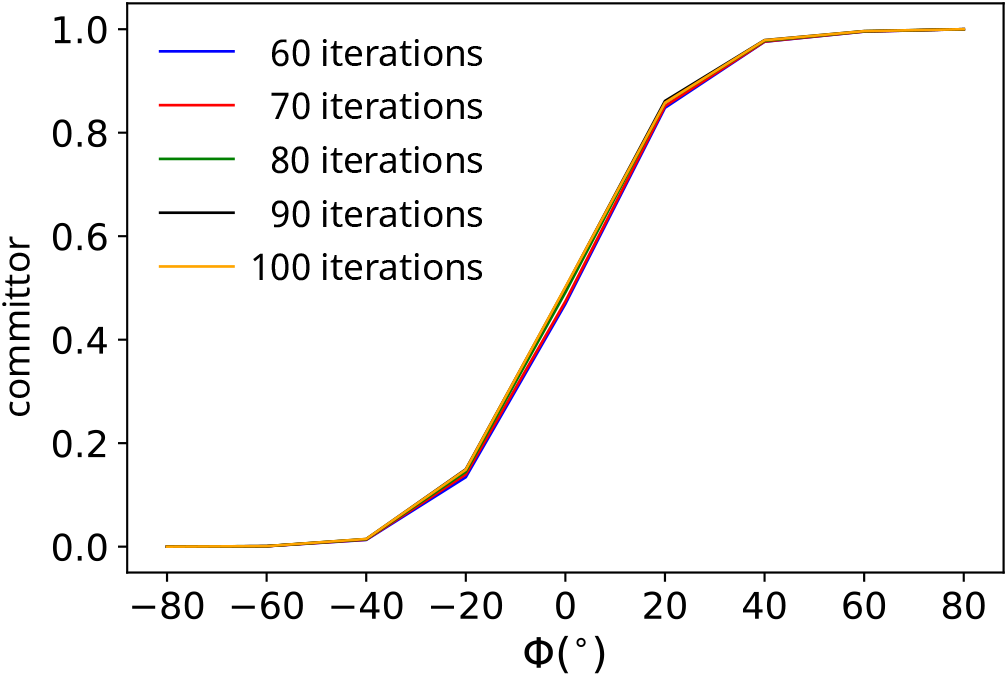
Committor function along the *ϕ* coordinate from M-WEM simulation.

Next, run the `free-energy-landscape.ipynb` to reconstruct the free energy landscape along Phi and Psi coordinates from the M-WEM data. It will first produce the unscaled probability distribution, rescale it and then compute the rescaled free energy landscape. The final free energy landscape should look like **Figure 12**.

**Figure 12.**
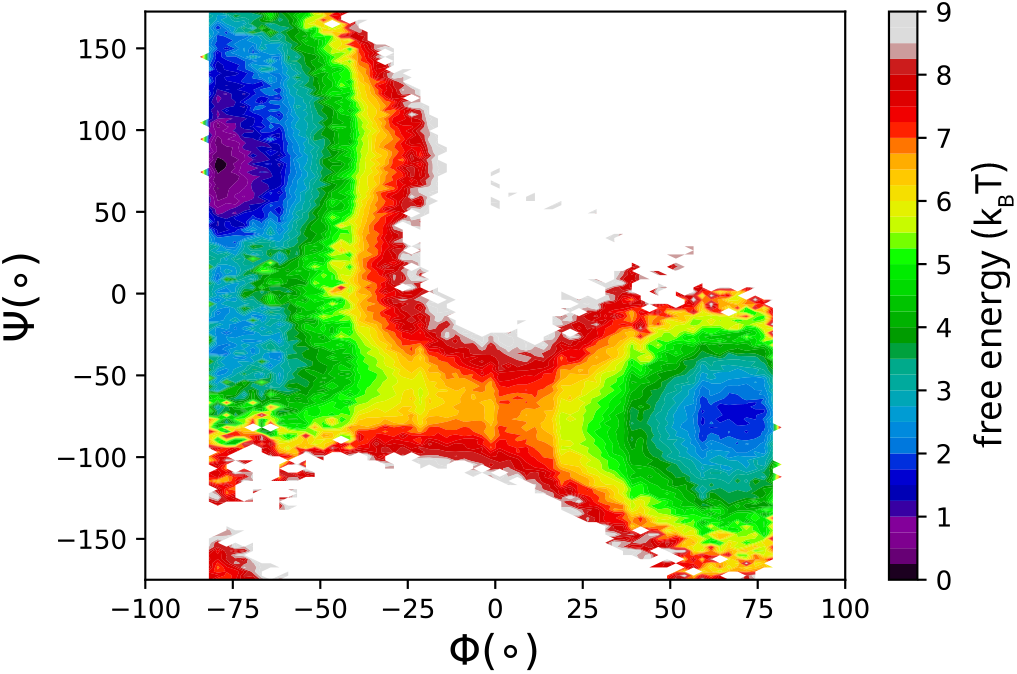
Free energy landscape reconstructed from an M-WEM simulation.

**Figure 13.**
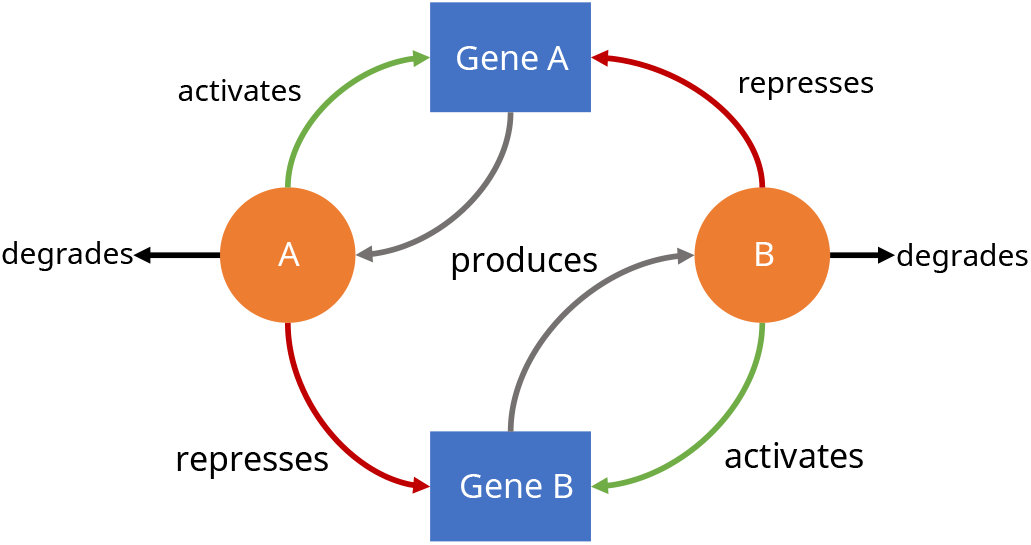
Two-gene network of exclusive mutual inhibition and self activation.

**Figure 14.**
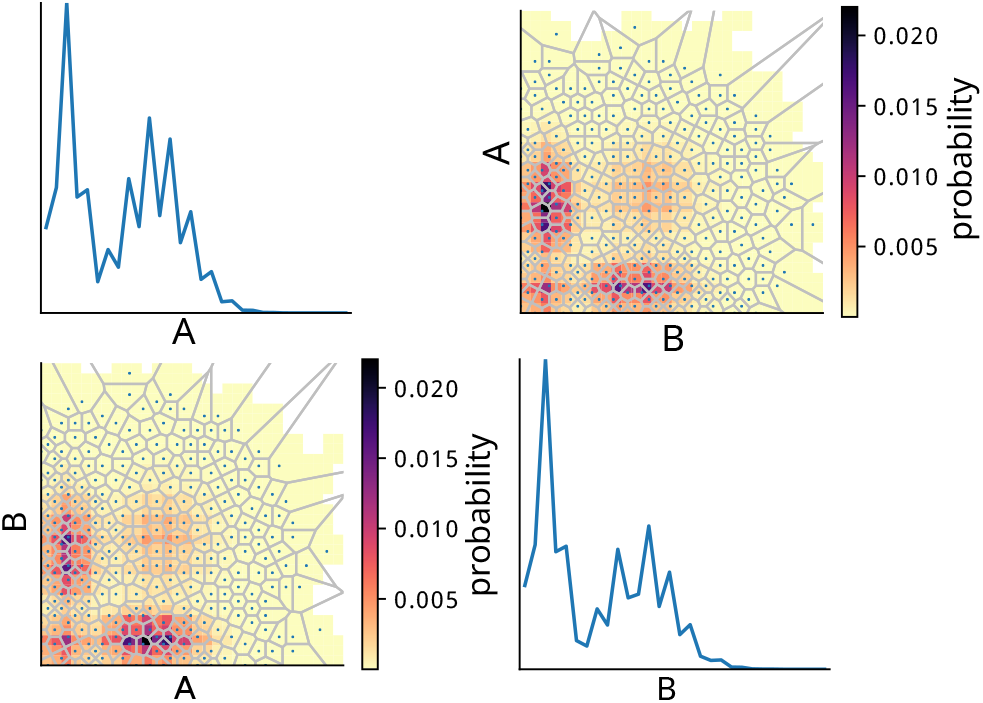
Probability distributions as a function of the two-dimensional progress coordinate (off-diagonal) and each dimension of the progress coordinate (diagonal). Grey lines in each off-diagonal plot delineate adaptive Voronoi bins used during the WE simulation.

**Figure 15.**
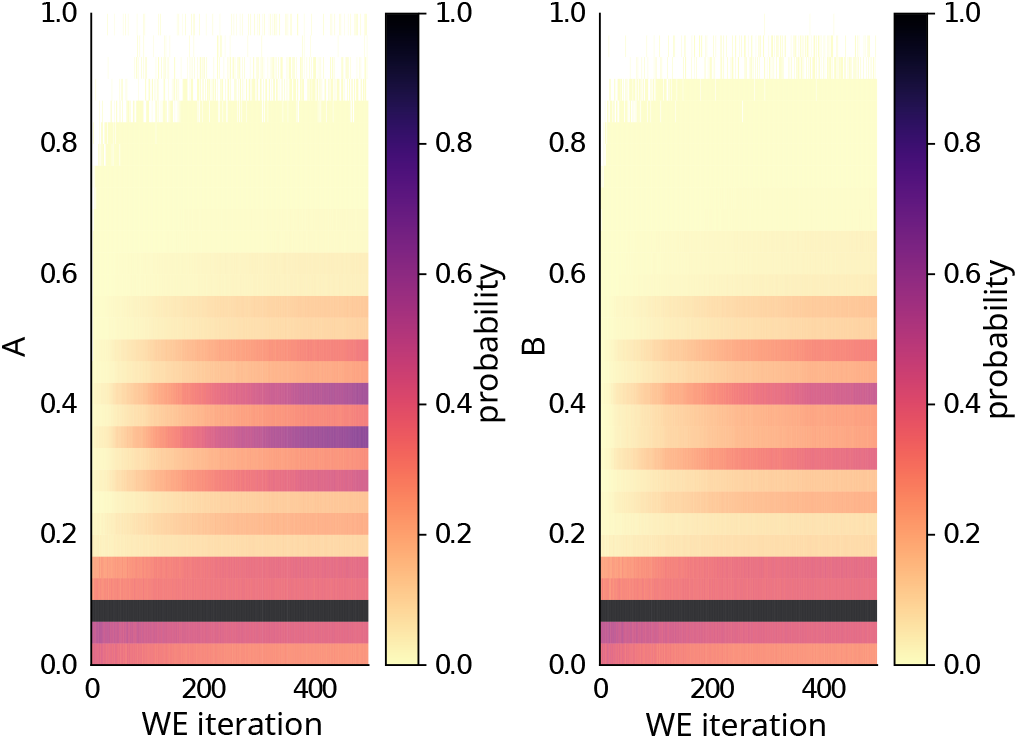
Probability distributions of each observable (A or B) of the two-dimensional progress coordinate as a function of WE iteration. The striped nature of the distributions is due to the fact that A and B are discrete as opposed to continuous observables.

Note that before executing any notebook, you will need to set the kernel to the environment in which you installed the M-WEM software. If the kernel is not available, activate the Jupyter notebook for that environment by executing:

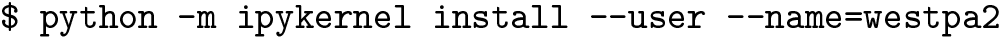

Replace westpa2 with the environment in which you installed M-WEM.

#### 3.5.7 Conclusion

This tutorial presents the Markovian Weighted Ensemble Milestoning (M-WEM) Python package for use with the WESTPA 2.0 software package to estimate equilibrium and non-equilibrium observables for the alanine dipeptide. In the M-WEM approach, the WE strategy is applied to enhance the efficiency of the Markovian milestoning approach to accelerate the convergence between milestones. While it is not possible to use this approach to generate continuous pathways between the initial and final states of a rare-event process, the M-WEM approach can be highly efficient in the calculation of “end-point properties” such as the MFPT and free energy differences between the two states. Beyond the alanine dipeptide, the M-WEM approach has been applied to more complex processes such as receptor-ligand binding, yielding the *k_on_*, *k_off_*, and binding free energy for the trypsin benzamidine complex [10].

### 3.6 Systems Biology Simulations using the WESTPA/BNG Plugin

#### 3.6.1 Introduction

This tutorial focuses on a scenario in systems biology in which the WE strategy can be useful: enhanced sampling of rare events in a non-spatial model. Here we focus on a BioNetGen language (BNGL) rule-based model for a biological signaling network that consists of a set of structured molecule types and a set of rules that define the interactions between the molecule types. While the average steady-state behavior of the model can be obtained using ordinary differential equations, the full kinetics of the model can only be obtained from stochastic simulations. However, adequate sampling of any rare events in the model can be a challenge for stochastic simulations. In this tutorial, we will use WESTPA to orchestrate BNGL simulations that are propagated by the BNG software package. As mentioned above, WESTPA is interoperable with any stochastic dynamics engine, including the BNG software.

##### Learning objectives

We will simulate a BNGL rule-based model of a two-gene switch motif that exhibits mutually exclusive activation and inhibition. Specific learning objectives include:

1. How to install the WESTPA/BNG plugin and set up a WESTPA/BNG simulation;
2. How to apply adaptive Voronoi binning, which can be used for both non-spatial and molecular systems;
3. How to run basic analyses tailored for high-dimensional WESTPA/BNG simulations.

#### 3.6.2 Prerequisites

Users should have a working knowledge of BNGL models (http://bionetgen.org) and the WESTPA 2.0 software package. This tutorial will make use of the WEBNG Python package, which facilitates the integration of WESTPA with the BNG software and requires Python 3.7 or later versions. To install the WEBNG package:

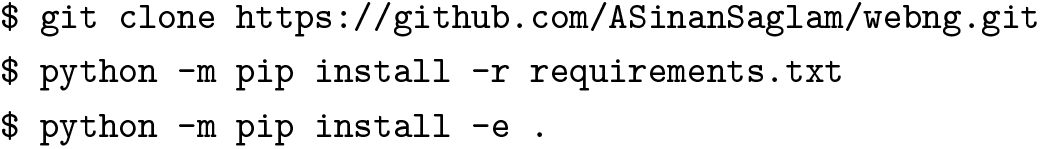

For common installation issues, see https://webng.readthedocs.io/en/latest/quickstart.html#installation. Alternatively the user can use a Docker container where the environment is already prepared correctly using the command:

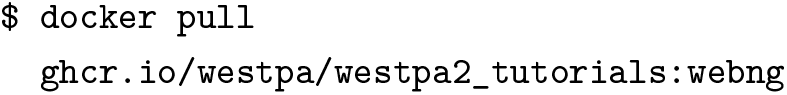

Note that this requires a Docker installation, for more information see Docker documentation (https://docs.docker.com/get-docker). Once the docker image is downloaded, you can run the image with:

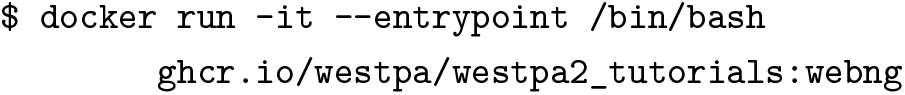

##### Computational requirements

This tutorial requires ~500 MB disk space. The simulation takes at most 1 hour of wall-clock time using a single CPU core of a 3.2GHz Intel Core i7 processor. We recommend using the `WEBNG` package on a Unix system. While the package has not been tested on Windows systems, one can try using the Windows subsystem for Linux (WSL; https://docs.microsoft.com/en-us/windows/wsl/install).

#### 3.6.3 Setting up the simulation

##### The model

Our BNGL model consists of two genes, gene A and gene B, that are transcribed to produce proteins A and B, respectively. Protein A binds to the gene A promoter site to activate protein A production and to the gene B promoter site to repress B production. Likewise, protein B activates gene B and represses gene A. The two most populated states therefore consist of either (1) high quantities of protein A and low quantities of protein B, or (2) high quantities of protein B and low quantities of protein A. Transitions between these two states are rare events.

##### Preparing the simulation environment

For this portion of the tutorial, you can use either your own BNGL model or the ExMISA model described above, which is the default WEBNG example. WEBNG uses a YAML configuration file to set up a WESTPA folder. The `WEBNG template` subcommand gives you a YAML config file with the same defaults which you can then edit and use to generate the WESTPA simulation folder.

If you are using the default example, the command to generate the template is the following:

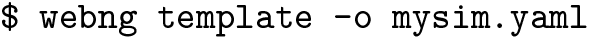

If you are using your own model file called `exmisa.bngl`, the command is:

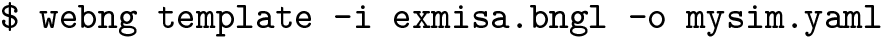

This command will generate the YAML file, `mysim.yaml`. For the full set of configuration options, see https://webng.readthedocs.io/en/latest/config.html. Path options are specified automatically using the libraries that are already installed. Propagator options will also be automatically populated according to the BNGL model. By default, this simulation setup will use an adaptive Voronoi binning scheme [15] due to the fact that rectilinear binning is not feasible for high-dimensional BNG models. The `center_freq` option sets the frequency of Voronoi bin addition, in units of WE iterations, `max_centers` is the maximum number of Voronoi bins that will be added, `traj_per_bin` is the number of trajectory walkers per Voronoi bin, and `max_iter` option sets the maximum number of WE iterations. All of these options can be modified after the simulation folder is set up (see https://github.com/westpa/westpa/wiki/User-Guide#Setting_Up_a_WESTPA_Simulation).

By default, the stochastic simulator is set to libroadrunner (http://libroadrunner.org). To use this simulator, we must first convert the BNGL model to a systems biology markup language (SBML) model. Next, we use the WEBNG software to compile the SBML model into a Python object, which allows for efficient simulation of the model. WEBNG also supports the use of the `BNG` simulation package. However, the use of this package will result in higher file I/O operations. Any other stochastic simulator will require the use of a custom WESTPA propagator.

#### 3.6.4 Running the WE simulation

To run the simulation, we first need to generate a WESTPA folder using the YAML configuration file generated in the previous step:

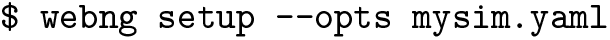

The above command will use the path option `sim_name` as the WESTPA folder, which is automatically set to the model name in the folder you ran the template command. Next, we initialize the simulation and run the model in a serial mode using a single CPU:

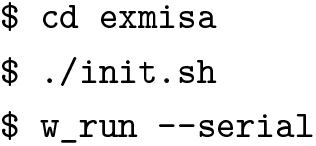

To run the model in a parallel mode using multiple CPU cores, please refer to WESTPA documentation for options available with the `w_run` command-line tool. The resulting simulation can be found in the `exmisa/` folder directory.

#### 3.6.5 Analyzing the WE simulation

To analyze the simulation, we will use the WEBNG package. To begin, we edit the YAML file under the folder that contains the configuration file `mysim.yaml`, setting `analyses.enable`, `analyses.average.enable`, and `analyses.evolution.enable` to `True`; and `analyses.average.first-iter` to the simulation half point (default: 50). To run the analysis, we use the following command:

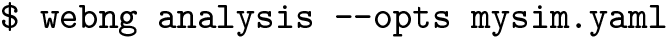

The above command will generate an `analysis/` folder in the simulation folder, run the analyses, and generate associated figures.

By default, `average.png` provides an N×N matrix of plots of the average two-dimensional probability distributions of each observable (dimension) of the WE progress coordinate and each of the other observables. The `evolution.png` file gives the time-evolution (number of WE iterations) of probability distributions for each observable and can be used to assess the convergence of simulation, making modifications to the binning scheme if necessary.

The average two-dimensional probability distributions reveal a total of four states: a low A/low B state, the symmetric low A/high B state and high A/low B states, and a high A/high B state. The fourth state is the least probable while the first three states are all highly probable. Transitions from low A/high B to high A/low B states are difficult to sample and transitions from low A/high B to high A/high B states are even more difficult to sample. The WE algorithm allows the user to sample these states and transitions between the states. All analyses should be based on the portion of the simulation that is done evolving. If the simulation is still evolving, we recommend extending the simulation until the observables of interest are reasonably converged.

#### 3.6.6 Conclusion

As demonstrated by this tutorial, the WEBNG Python package provides a framework for applying the WESTPA 2.0 software package to BNGL models with minimal user input and simplified installation. The adaptive Voronoi binning scheme enables efficient application of high-dimensional progress coordinates for both molecular and non-spatial systems. Voronoi bins can be effective for exploratory simulations, placing bins as far away as possible from previous bins to inform the creation of a custom binning scheme for sampling the rare-event process of interest. However, such bins may not be as effective for surmounting barriers (e.g., compared to the MAB scheme [18]), as demonstrated by the probability distribution as a function of the WE progress coordinate where many bins near the edges of the configurational space are occupied, but are not of interest. Future work with WEBNG will include more detailed analysis options such as automated clustering, generation of networks from bins and clusters, and the estimation of rate constants for transitions between the clusters.

## 4 Author Contributions

AT Bogetti, JMG Leung, JD Russo, S Zhang, JP Thompson, AS Saglam, and D Ray developed and wrote the tutorials; the primary author of each tutorial is designated as a co-first author of the overall manuscript. LT Chong, DM Zuckerman, DN LeBard, MC Zwier, JL Adelman, I Andricioaei, and JR Faeder provided guidance for tutorial development. AT Bogetti, JMG Leung, LT Chong, DM Zuckerman, and MC Zwier wrote the introductory sections leading up to the tutorials. RC Abraham helped generate trajectory data for **Advanced Tutorial 3.1**.

## 5 Other Contributions

We thank the many users of the WESTPA software who have provided important feedback for improving our tutorials and documentation over the years.

## 6 Potentially Conflicting Interests

The authors declare the following competing financial interest(s): L.T.C. is a current member of the Scientific Advisory Board of OpenEye Scientific and an Open Science Fellow with Roivant Sciences. S Zhang, JP Thompson, and DN LeBard are employees of OpenEye Scientific.

## 8 Content and Links

All files needed for the tutorials can be found at https://github.com/westpa/westpa2_tutorials.

